# Sleep spindles, ripples, and interictal epileptiform discharges in the human anterior and mediodorsal thalamus

**DOI:** 10.1101/2021.08.12.456047

**Authors:** Orsolya Szalárdy, Péter Simor, Péter Ujma, Zsófia Jordán, László Halász, Loránd Erőss, Dániel Fabó, Róbert Bódizs

**Affiliations:** Institute of Behavioural Sciences, Faculty of Medicine, Semmelweis University; National Institute of Clinical Neurosciences; Institute of Cognitive Neuroscience and Psychology, RCNS; Eötvös Loránd Unniversity, Faculty of Education and Psychology

**Keywords:** sleep spindle, interictal epileptic discharges, general intelligence, thalamus, epilepsy

## Abstract

Sleep spindles are major oscillatory components of Non-Rapid Eye Movement (NREM) sleep, reflecting hyperpolarization-rebound sequences of thalamocortical neurons, the inhibition of which is caused by the NREM-dependent activation of GABAergic neurons in the reticular thalamic nucleus. Reports suggest a link between sleep spindles and several forms of interictal epileptic discharges (IEDs) which are considered as expressions of pathological off-line neural plasticity in the central nervous system. Here we investigated the relationship between thalamic sleep spindles, IEDs and ripples in the anterior and mediodorsal nuclei (ANT and MD) of epilepsy patients. Whole-night LFP from the ANT and MD were co-registered with scalp EEG/polysomnography by using externalized leads in 15 epilepsy patients undergoing Deep Brain Stimulation protocol. Slow (∼12 Hz) and fast (∼14 Hz) sleep spindles were present in the human ANT and MD. Roughly, one third of thalamic sleep spindles were associated with IEDs or ripples. Both IED- and ripple-associated spindles were longer than pure spindles. IED-associated thalamic sleep spindles were characterized by broadband increase in thalamic and cortical activity, both below and above the spindle frequency range, whereas ripple-associated thalamic spindles exceeded pure spindles in terms of 80–200 Hz thalamic, but not cortical activity as indicated by time-frequency analysis. These result show that thalamic spindles coupled with IEDs are reflected at the scalp slow and beta-gamma oscillation as well. IED density during sleep spindles in the MD, but not in the ANT was identified as correlates of years spent with epilepsy, whereas no signs of pathological processes were correlated with measures of ripple and spindle association. Furthermore, the density of ripple-associated sleep spindles in the ANT showed a positive correlation with general intelligence. Our findings indicate the complex and multifaceted role of the human thalamus in sleep spindle-related physiological and pathological neural plasticity.

## Introduction

Higher order brain functions rely on distributed networks involving the cortex and subcortical regions. The thalamus plays a crucial role in the proper functioning of these networks: thalamo-cortical loops are associated with sensory processing, memory formation, executive functions, many of which are tightly related with sleep state-dependent neural oscillations. The thalamus also plays an important role in the coordinated connection between the cortex and hippocampus (Latchoumane et al., 2017). The human anterior thalamic nucleus (ANT) is a higher-order thalamic nucleus which is interconnected with the hippocampus, and plays an important role in episodic memory formation (Aggleton et al., 2010). It has reciprocal connections with the anterior cingulate cortex, retrosplenial cortex, and subiculum, and selective inactivation of the ANT leads to impaired memory formation similarly to hippocampal lesions. Furthermore clinical and experimental results suggest that in Wernicke-Korsakoff syndrome and in thalamic stroke, the episodic memory deficit is closely related to the dysfunction of the ANT (Child & Benarroch, 2013). Furthermore, the ANT plays an important role in the propagation of epileptic seizures and therefore became an important target for deep brain stimulation (DBS) which serves as a treatment for medically refractory epilepsy (Salanova, 2018). One other important higher-order nucleus in the human thalamus is the mediodorsal nucleus (MD) which is reciprocally connected with the medial prefrontal cortex (mPFC) and also receives inputs from parahippocampal regions, and therefore it is assumed to interact with the cortex and hippocampus in declarative memory formation during non-REM (NREM) sleep (Mitchell & Chakraborty, 2013). Reduced MD volume was associated with decreased sleep spindle density in frontal brain regions which suggest that the MD is involved in sleep spindle generation and/or propagation (Buchmann et al., 2014). In addition, the degeneration of MD in patients suffering from the prion disease fatal familial insomnia is associated with lack of sleep spindles (Lugaresi et al., 2006).

Sleep spindles are NREM sleep state-specific oscillations characterized by waxing/waning 10–16 Hz waveforms that are generated by the reticular nucleus of the thalamus and are propagated to other brain regions by thalamo-cortical circuits (Huguenard & McCormick, 2007; Steriade, 2005). Animal studies confirmed that dynamic reticular thalamic-thalamo-cortical interactions are responsible for the generation of sleep spindles (Steriade, 2005; Steriade et al., 1987), which were associated with cognitive functions such as learning and memory consolidation (Cairney et al., 2018; Saletin et al., 2011), and intellectual ability (Fogel & Smith, 2011; Ujma, 2018a; Ujma et al., 2020). Sleep spindles tend to co-occur with and coordinate the faster oscillations, called hippocampal ripples (∼80–140 Hz), providing efficient off-line plasticity windows for memory consolidation and reorganization (Girardeau & Zugaro, 2011). Besides neural plasticity sleep spindles were associated with pathological off-line plasticity in epileptic models and human epilepsy patients (Gelinas et al., 2016). Findings regarding the strong relationship between NREM sleep oscillations, neuroplasticity and epilepsy indicate that sleep-associated interictal epileptic discharges (IEDs) harm cognitive functions, inducing a significant cognitive loss (Halász et al., 2019). Whereas the reticular nucleus of the thalamus is responsible for spindle generation, higher order thalamic nuclei are also involved in declarative memory processes (Van Der Werf et al., 2003), and general intellectual ability (Burgaleta et al., 2014). In spite of the large number of animal studies investigating the role of the thalamus in memory formation and higher level cognitive function, the direct measurements of the activity of these thalamic nuclei within the human brain are surprisingly rare.

The aim of the current study was to assess the function of ANT and MD in sleep-related neural plasticity and intellectual ability (IQ) measured by the association between sleep spindles, thalamic ripples and IEDs during NREM sleep. It was proposed that sleep-related epileptic transformation of normal neurological networks, involving the hippocampus, thalamus and cortex, interfere with sleep-related synaptic homeostasis, neural plasticity, and cognitive functioning (for a recent review, see Halász & Szűcs, 2020). In this pathway, the role of the human ANT and MD is not clear. Although, different aspects of sleep spindle-related dynamic thalamocortical interactions were revealed, relationships with epilepsy, IEDs and ripples in the thalamus were not analyzed. A recent report (Rektor et al., 2016) indicated the occurrence of high frequency oscillations (HFOs) such as ripples in the human thalamus, however the number of results confirming the involvement of HFOs in the thalamus is remarkably sparse. In that study, HFOs in the ripple frequency band (up to 240 Hz) were identified in the ANT suggesting the involvement of thalamic ripples in pathological processes related to epilepsy. However, no direct connection was revealed between epilepsy and thalamic ripples. In contrast, IEDs in the human thalamus, such as in the ANT LFP records were observed and suggested the involvement of ANT in the propagation of epileptic activity and to contribute to the epileptic network (Hodaie et al., 2002; Sweeney-Reed et al., 2016). In several former studies, sleep spindles were recorded from the human thalamic nuclei, including the ANT and the posterior thalamus/medial pulvinar nucleus of epilepsy patients undergoing DBS or invasive presurgical neurophysiological investigation (Bastuji et al., 2020; Mak-Mccully et al., 2017; Tsai et al., 2010). Here we investigated the association between thalamic sleep spindles, interictal epileptiform discharges, ripples, and their associations with intellectual ability within two higher-order thalamic nuclei in human subjects, the ANT and the MD.

## Methods

### Participants

Subjects were 15 pharmacoresistant, surgically non-treatable epilepsy patients (M_age_ = 36.9 years, range: 17–64; 7 female) participating in the ANT deep brain stimulation (DBS) protocol at the National Institute of Clinical Neurosciences, Budapest, Hungary. Clinical and demographic data are reported in Table 1. The research was approved by the ethical committee of the National Institute of Clinical Neurosciences. Patients signed informed consent for participating in the study. General intelligence (IQ) was measured prior to the surgery by Wechsler Adult Intelligence Scales, 4th edition IQ from 11 patients who underwent on neuropsychological testing in the context of their clinical routine presurgical assessment.

**Table 1.**
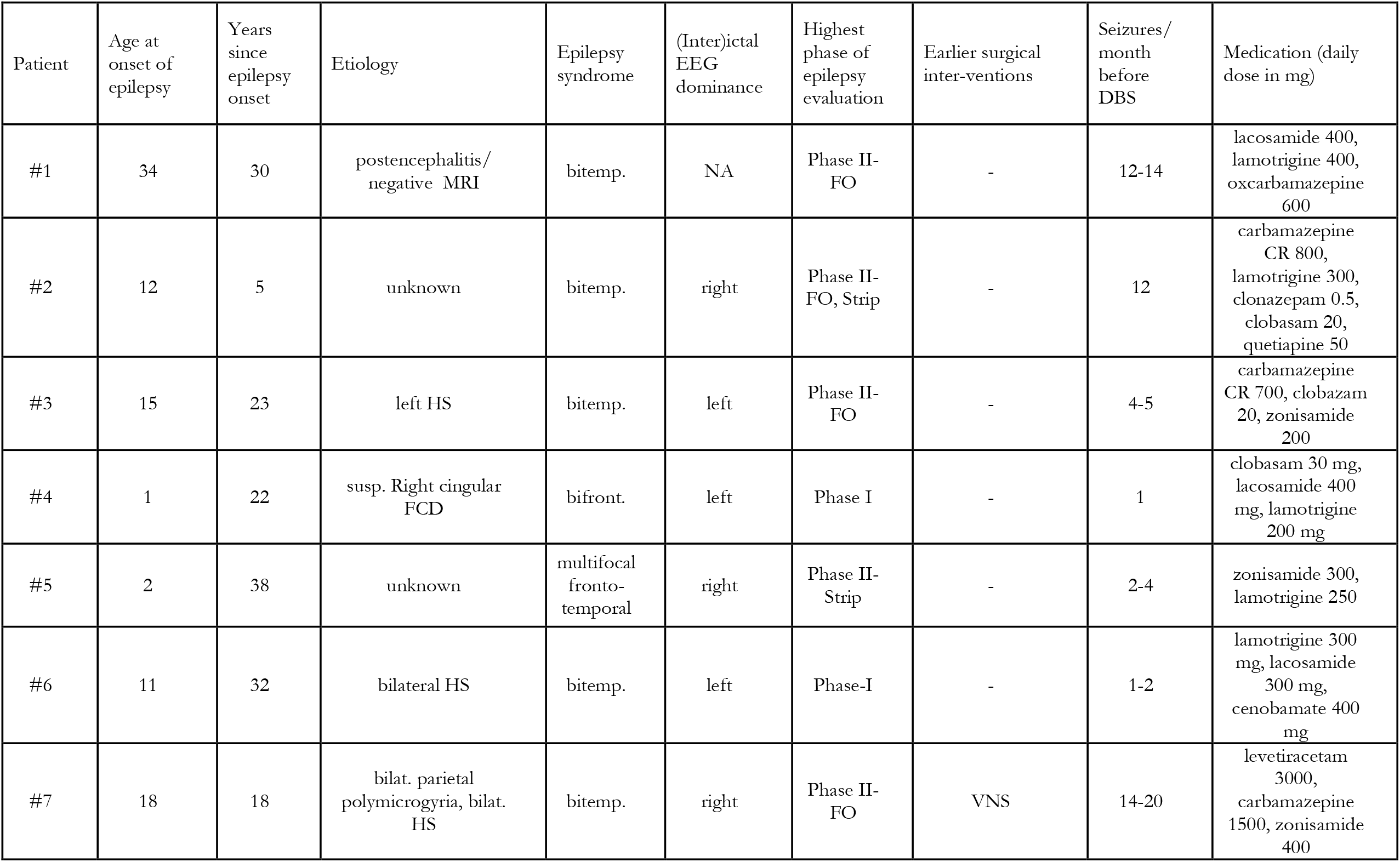
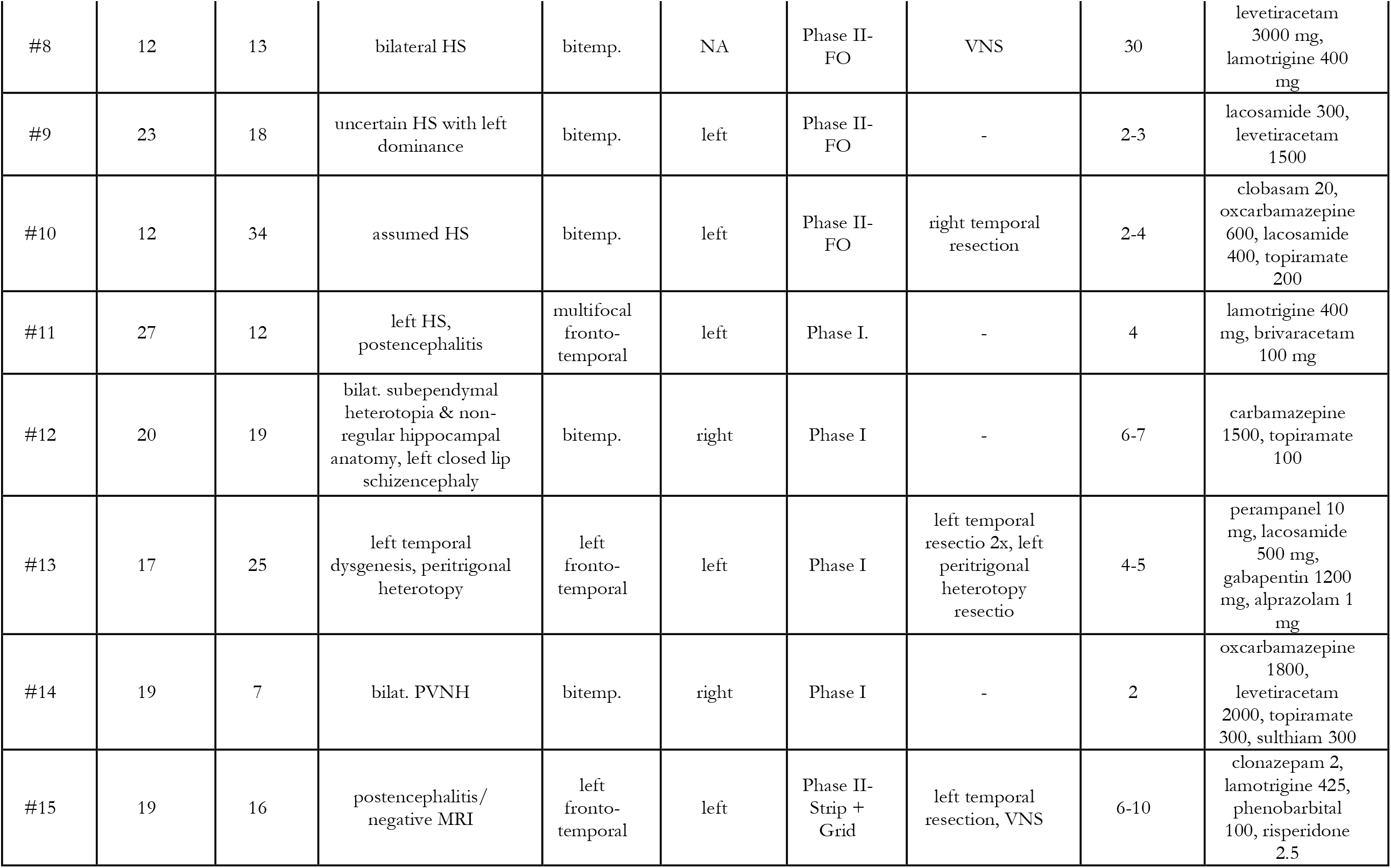
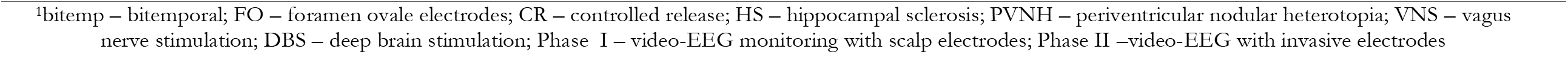
Clinical and demographic data of the patients

### Procedure

A pair of quadripolar Medtronic DBS electrodes were bilaterally and stereotaxically inserted into the anterior part of the thalamus during general anaesthesia. Contact lengths of the electrode are 1.5 mm, whereas intercontact spacing is 0.5 mm. Electrode diameter is 1.27 mm. Besides frontal transventricular and extraventricular trajectories, the posterior parietal extraventricular approaches were used in accordance with the decisions of the clinical-neurosurgical team. In accordance with the clinical protocol patients underwent a 48 hour, postsurgical video-EEG monitoring with externalized thalamic leads, that is thalamic LFP co-registered with scalp EEG/polygraphy. The first 24 hours were recorded without thalamic stimulation (DBS-off), whereas the effects of DBS were tested in the second 24 hours period (DBS-on). The whole-night records of the DBS-off records were analysed in the present study.

### Individual localization of thalamic contacts

Thalamic contacts were localized by using the procedure described below (see also Simor et al., 2021). In order to individually localize the thalamic contacts preoperative MRI and postoperative CT images were co-registered using tools available in the FMRIB Software Library (FSL, Oxford, FLIRT, linear registration, 6 degrees of freedom). Threshold was applied on the co-registered CT scans to achieve the desired level of density for proper identification of the lead, thus removing the surrounding brain tissue. Coordinates of the most distal point of the lead were identified and a more proximal point was selected along the line of the contacts to mathematically reconstruct the coordinates of the center point of each contact using Euclidean distance in three dimensional space. These points superimposed over the T1 MRI image provided a guideline for contact localization by examining their location to the anatomical boundaries of the ANT (See Figure 1). Anatomical positions according to standard coordinates of the contacts were double-checked by using the mamillothalamic tract as an anatomical guide to localize the ANT. In case of convergent results of these two methods, the ANT (for 15 patients) and MD (for 10 patients) contacts were considered as subjects for further digital signal processing in the present study.

**Figure 1.**
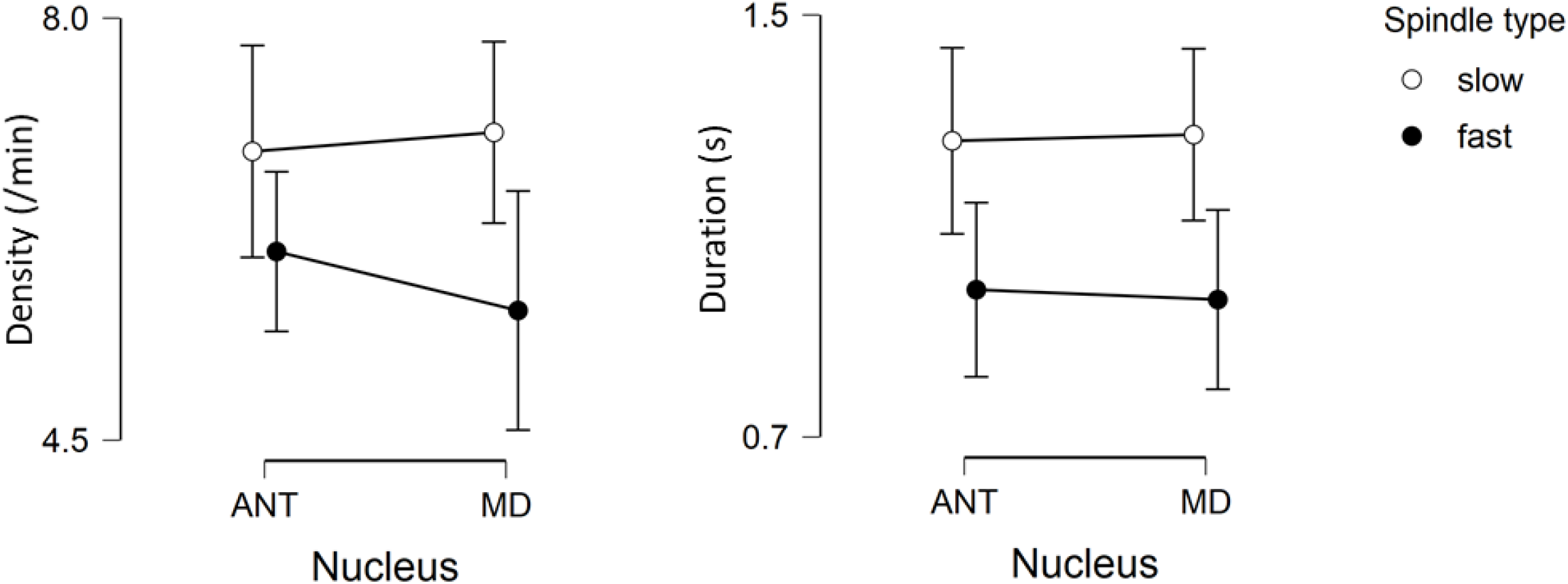
Overall sleep spindle density (left panel) and duration (right panel) in the ANT (left) and MD (right), separately for the slow spindles (white dots) and fast spindles (black dots).

### EEG and LFP data recording a preprocessing

ANT, MDT (LFP) and all-night sleep EEG signals were recorded by SD-LTM 64 Express EEG/polygraphic recording system. Physiological signals were recorded at 8192 Hz/channel effective sampling rate with 22 bit precision and hardware input filters set at 0.02 (high pass: 40 dB/decade) and 450 Hz (low pass: 40 dB/decade). Data was decimated by a factor of 4 by the firmware resulting in stored time series digitized at 2048 Hz/channel. LFP signals were assessed by bilateral (L – left, R – right) quadripolar electrodes applying bipolar reference scheme and focusing only on those leads which were derived from two adjacent contacts positioned in the same thalamic nucleus or alternatively from the BRIDGE area (bipolar recordings with one contact within the ANT and the second contact in adjacent tissue Deutschová et al., 2021). The number of ANT and MD derivations recorded for each patient is reported in Table 2. Scalp EEG was recorded according to the international 10-20 system (Fp1, Fp2, Fpz, F3, F4, F7, F8, C3, C4, T3, T4, T5, T6, P3, P4, O1, O2, Oz; Jasper, 1958) extended with the inferior temporal chain (F9, F10, T9, T10, P9, P10) and two anterior zygomatic electrodes (ZA1, ZA2; Manzano et al., 1986), with the reference placed at the CP1 and ground placed at CP2 locations. Submental electromyograms (EMGs) were recorded by bipolarly referenced electrodes placed on the chin. For two patients (#1 and #2) the EEG electrode set up were slightly different from the above described montage. For patient #1 no Fpz, P9, P10 and EMG time series were recorded, and for patient #2 F3, F4, C3, C4, P3 and P4 electrodes were missing, although the electrode position at Pz was present in this patient. Data from missing electrodes were treated as missing values in further analyses.

**Table 2.** Number of ANT and MD derivations for each patient

| **Patient** | **Total** | **Left ANT** | **Right ANT** | **Total** | **Left MD** | **Right MD** |
| --- | --- | --- | --- | --- | --- | --- |
|  | **number of**<br>**ANT**<br>**derivations** |  |  | **number of**<br>**MD**<br>**derivations** |  |  |
| #1 | **1** | 0 | 1 | **1** | 0 | 1 |
| #2 | **2** | 0 | 2 | **1** | 0 | 1 |
| #3 | **2** | 0 | 2 | **1** | 0 | 1 |
| #4 | **3** | 2 | 1 | **0** | 0 | 0 |
| #5 | **1** | 1 | 0 | **1** | 1 | 0 |
| #6 | **4** | 2 | 2 | **0** | 0 | 0 |
| #7 | **1** | 0 | 1 | **2** | 0 | 2 |
| #8 | **6** | 3 | 3 | **0** | 0 | 0 |
| #9 | **3** | 1 | 2 | **1** | 0 | 1 |
| #10 | **3** | 2 | 1 | **2** | 1 | 1 |
| #11 | **3** | 2 | 1 | **0** | 0 | 0 |
| #12 | **2** | 1 | 1 | **2** | 1 | 1 |
| #13 | **2** | 1 | 1 | **1** | 1 | 0 |
| #14 | **4** | 2 | 2 | **0** | 0 | 0 |
| #15 | **1** | 0 | 1 | **1** | 0 | 1 |

Continuous EEG and LFP recordings were automatically segmented into 90 minutes chunks by the recording software. During the recording session there were no explicit light off and light on time, thus, the first and the last chunk for data analysis were selected as follows: a 90 minutes segment were required to contain at least 10 minutes of continuous sleep in the second or in the first halves, respectively for the first and last chunks. EEG data were offline re-referenced to the mathematically-linked T9 and T10 electrodes. All-night sleep records were scored for sleep-waking states and stages according to standard AASM criteria on 20 seconds basis (Berry et al., 2015) by an expert. Furthermore, artefactual segments were marked on 4 seconds basis and excluded from quantitative EEG and time series analyses.

### Sleep spindle detection and analysis

Non-artifactual NREM sleep EEG and thalamogram records were subjected to the Individual Adjustment Method (IAM) of sleep spindle analysis (Bódizs et al., 2009; Ujma, Gombos, et al., 2015). Frontally dominant slow and parietally dominant fast sleep spindles were defined for each patient and each EEG and thalamic channel using individual-specific frequency criteria, as well as associated individual- and derivation-specific amplitude criteria in the 9–16 Hz all-night NREM sleep. Thalamic sleep spindles were categorized on the basis of their association with IEDs and ripples. If one or more IED occurred between the onset and offset of a sleep spindle, the spindle was categorized as IED-associated spindle, referred SP(IED) along the text. If one or more ripple was detected during the spindle in the absence of IEDs, that spindle was categorized as ripple-associated spindle, SP(ripple). Spindles in which both IEDs and ripples were detected, were regarded as SP(IED). Furthermore, spindles without any association with IEDs and ripples were categorized as pure sleep spindles, SP(pure). The spindle densities (number per minutes) were calculated separately for each category. Sleep spindle density data were averaged across appropriate derivations within a nucleus, separately for the ANT and MD, and for the fast and slow spindles. Absolute numbers and spindle densities are summarized in Supplementary Table 3. The duration of sleep spindles was measured as the time interval from spindle onset and offset, separately for each spindle category. Spindle duration data from all derivation corresponding to same nucleus were averaged, separately for the ANT and MD, and for the fast and slow spindles.

### Detection and analysis of thalamic IEDs and ripples

Data analyses were performed by MATLAB version 9.5 (R2018b, The MathWorks, Inc., Natick, MA; https://www.mathworks.com/products/new_products/release2018b.html) using the EEGlab toolbox 14.1.2b. For removing potential electric power-related noise, 4 Hz bandstop filter was applied centered at 50 Hz and its harmonics up to 400 Hz (FIR filter, filter order: 3380). IED and ripple detection was performed on the basis of amplitude and frequency criteria within the NREM stages 2 and 3 sleep. Whole-night ANT and MD LFP signals were filtered with a 5 Hz highpass finite impulse response (FIR) filter (filter order: 3380). Thalamic IEDs were defined on the filtered signal by the absolute maximum of the filtered LFP signal between the time points the amplitude exceeded the mean NREM (stage 2 and 3) signal by 5 standard deviations (SD), separately for each derivation (for similar detection criteria, see Ujma, Simor, et al., 2015). Furthermore, if a consecutive IED were detected within 100 ms, the one with the largest amplitude was detected only. For thalamic ripple detection bandpass filtering of the whole-night ANT and MD LFP signals were performed at 80–200 Hz (FIR filter, transition bandwidth: filter order: 3380). Ripples were defined by the absolute maximum amplitude of the filtered LFP signal between the time points the amplitude exceeded by 4 SD the mean of the artefact-free NREM (stages 2 and 3) thalamograms. Elimination of consecutive ripples were performed within 55 ms (corresponding to the cycle length of the 18 Hz oscillation), choosing the one with the largest absolute amplitude. Density of the IEDs and ripples (IED and ripple number per minutes) were calculated for the whole non-artifactual NREM sleep (in stage 2 and 3; defined as IED(NREM) and ripple(NREM) density) and for sleep spindles (i.e. the density of IED or ripple events occurring during sleep spindle periods; defined as IED(sp) and ripple(sp) density, respectively). IED(NREM) and ripple(NREM) density and IED(sp) and ripple(sp) density were compared by two-tailed Student’s t-tests, separately for the ANT and MD. Mean thalamic and scalp IED density were also assessed by manual IED detection in ANT and MD LFP signals, based on approximately 20 minutes of continuous NREM sleep (stage 2 and/or 3), by two skilled experts in epileptology and clinical neurophysiology. Manually detected IED number were then standardized according to IED number per minutes of NREM sleep.

In order to test the convergent validity of the automatic IED and ripple detections, outcomes of these measures were correlated with manual IED detections (both thalamus and scalp), as well as with clinical epilepsy characteristics (seizure/months, year since epilepsy onset, age) by calculating the Pearson coefficients. Furthermore, Pearson correlations between SP(IED/pure/ripple) density and the same epilepsy characteristics (age, seizure/months, year since epilepsy onset, and manually detected thalamic and scalp IED density) were also tested. The results are reported in the Supplementary Material.

Overall sleep spindle density and duration were compared across nuclei performing repeated-measures ANOVA with the factors spindle type (slow vs. fast) × nucleus (ANT vs. MD), where missing values were handled as missing data. Comparison of SP(IED/ripple/pure) density and duration were performed by repeated-measure ANOVA with the factors spindle type (slow vs. fast) × association (IED vs. ripple vs. pure), separately for the ANT and MD. Correlations of spindle density (overall sleep spindle density, and separate analysis for the SP(IED/ripple/pure) density) with general intelligence were tested by Pearson coefficients, separately for the ANT and MD, and for the slow and fast spindles. Relationship between scalp-detected, parietal fast sleep spindle density (average of recording locations P3 and P4, except Patient #2, where Pz was used) and general intelligence was also tested. Statistical analyses were performed by STATISTICA 13.1 and JASP 0.11.0.0.

### Time-frequency decomposition of sleep spindles

Time-frequency wavelet analysis was performed on 4600 ms long epochs extracted in the -2300 to 2300 ms latency range around sleep spindle onsets (separately for the slow and fast SP(IED/ripple/pure)) on all ANT and MD derivations. Furthermore, time-frequency analysis was also performed on frontal and parietal scalp records using the same time intervals as for the ANT and MD sleep spindles (scalp EEG activity around thalamic sleep spindles). Data from the F3 and F4 electrodes were analyzed for the frontal region, except for patient #15, where Fp1 was used instead of F3 due to the signal quality. Parietal electrodes were P3 and P4, except for patient #2, where Pz was used instead of P3 and P4 as these electrodes were missing for this patient. Data were averaged across the frontal and parietal electrodes, separately.

Time-frequency analysis was performed by Morlet wavelet transformation with linearly increasing cycle numbers from 4 cycles to 20 cycles for the 1–450 Hz frequency range using 0.5 Hz frequency resolution in the 1–40 Hz range, 1 Hz frequency resolution in the 41–80 Hz range and 5 Hz frequency resolution in the 81–450 Hz range. The baseline interval was set at -2000 ms to 0 ms before sleep spindle onset. Time-frequency power spectra were averaged across all derivations within the same nucleus, separately for the fast and slow SP(IED/ripple/pure). Statistical comparison of SP(ripple) vs. SP(pure) and SP(IED) vs. SP(pure) were performed by nonparametric randomization test separately for the slow and fast spindles and for the ANT, MD and scalp (frontal and parietal), applying the Monte-Carlo estimate of significance probabilities (1000 permutations), using the Fieldtrip toolbox (Oostenveld et al., 2011). The alpha level was set at 0.025, avoiding multiple comparisons. Bandwise comparisons were performed by averaging each time-frequency power spectra data point within the 300–800 ms latency range after spindle onset for the sigma (11–16 Hz) band. Statistical analysis of bandwise comparison was performed by repeated-measures ANOVA with the factors association SP(IED/ripple/pure) separately for spindle type (slow, fast) and scalp measurement (frontal and parietal).

## Results

### 1. Thalamic sleep spindles, ripples and IEDs

#### Overall sleep spindle density and duration in the ANT and MD

Although the main effect of spindle type was not significant (F(1,9) =3.689, p=0.087), nucleus and spindle type factors interacted in predicting higher slow over fast spindle density differences in MD as compared to ANT (Figure 1; F(1,9)=6.055, p=0.036, ηp2=0.402). Post-hoc test revealed a significant difference between the density of slow and fast sleep spindles within the same thalamic nuclei (ANT: p=0.007, MD: p<0.001), and also between the two nuclei (i.e. ANT slow spindle density was higher than MD fast spindles density, p=0.002) although, slow sleep spindle density was not different between the ANT and MD, similarly to fast spindle density (p=0.104, both). The duration of fast sleep spindles were significantly shorter compared to the slow sleep spindles which was confirmed by the main effect of spindle type (F(1,9)=5.440, p=0.045, ηp2=0.377). Supplementary Table 2. summarizes the absolute number, density and duration of slow and fast sleep spindles, separately for each patient, averaged across each derivation within the same nucleus.

#### IED and ripple density during sleep spindles and overall NREM sleep

IEDs were more frequently coupled with sleep spindles as compared to overall NREM sleep. That is, significantly higher IED(sp) density as compared to IED(NREM) density was measured both in the anterior (t=5.745, df = 14, p<0.001) and mediodorsal (t=3.992, df = 9, p=0.003) thalamic nuclei (Figure 2). Ripple(NREM) and ripple(sp) density were not significantly different from each other, neither in the ANT (t=1.334, df = 14, p=0.203) nor in the MD (t=1.986, df = 9, p=0.078). Thus, IEDs were more frequently coupled with sleep spindles than with other NREM segments.

**Figure 2.**
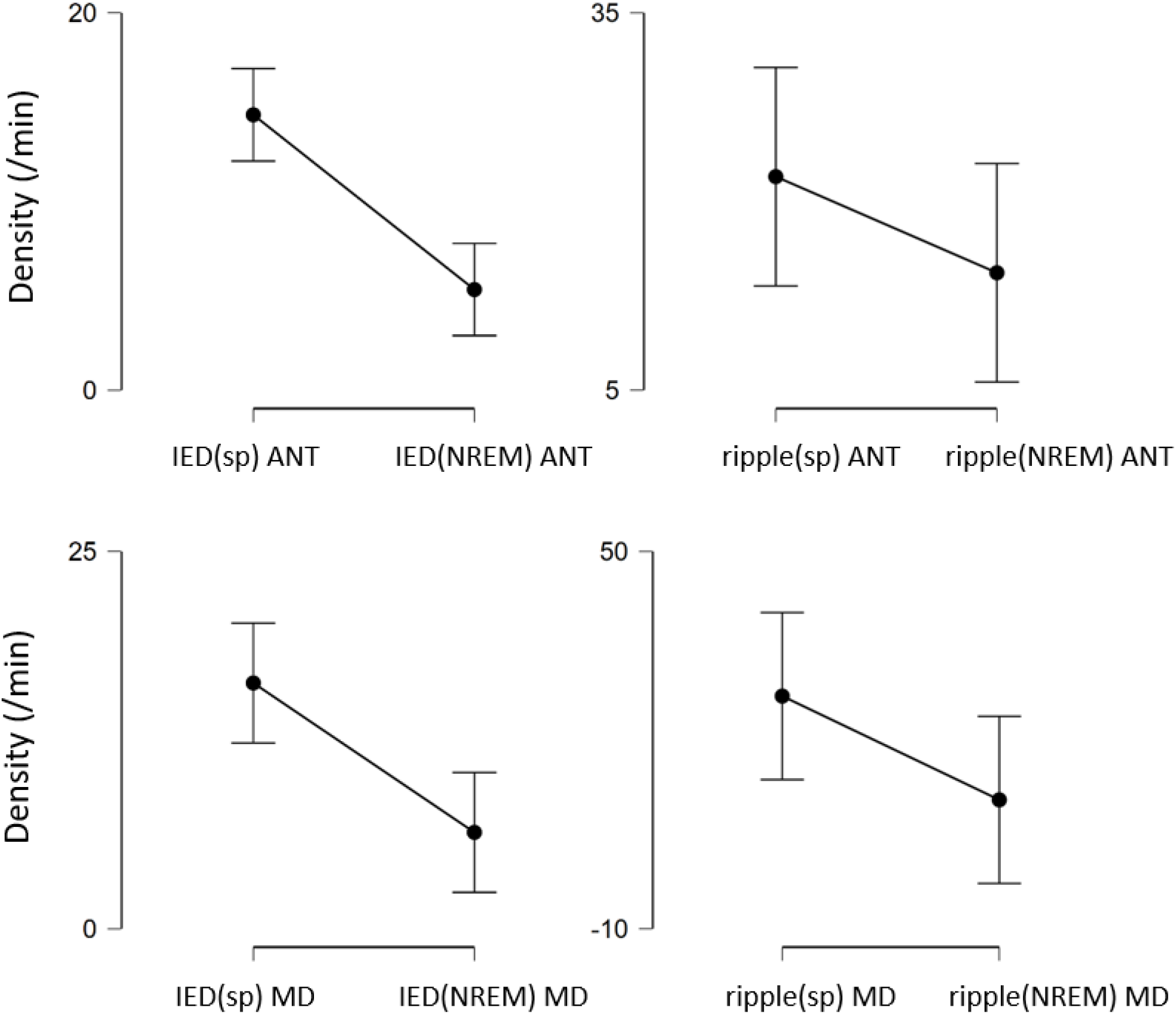
IED (left panels) and ripple (right panels) density in the ANT (top panels) and MD (bottom panels).

### 2. Sleep spindles associated with ripples and IEDs

#### Sleep spindle density and duration (SP(pure/IED/ripple) in the ANT and MD

Figure 3 shows examples for fast sleep spindles from the same patient and nucleus, associated with and without IEDs and ripples. On average 20.034% of sleep spindles were associated with IEDs, 10.847% with ripples, and 69.119% were pure sleep spindles. Significant main effect of co-occurrence was found for the spindle density, separately for the ANT and MD (Figure 4; F(2,28)=92.267, p<0.001, ε=0.745, η_p_^2^=0.868 for the ANT and F(2,18)= 34.008, p<0.001, ε=0.741, η_p_^2^= 0.791 for the MD), which resulted by the higher SP(pure) density compared to the SP(IED) and SP(ripple; p<0.001, all). No further main effect or interaction reached the critical alpha level measured for the density of sleep spindles.

**Figure 3.**
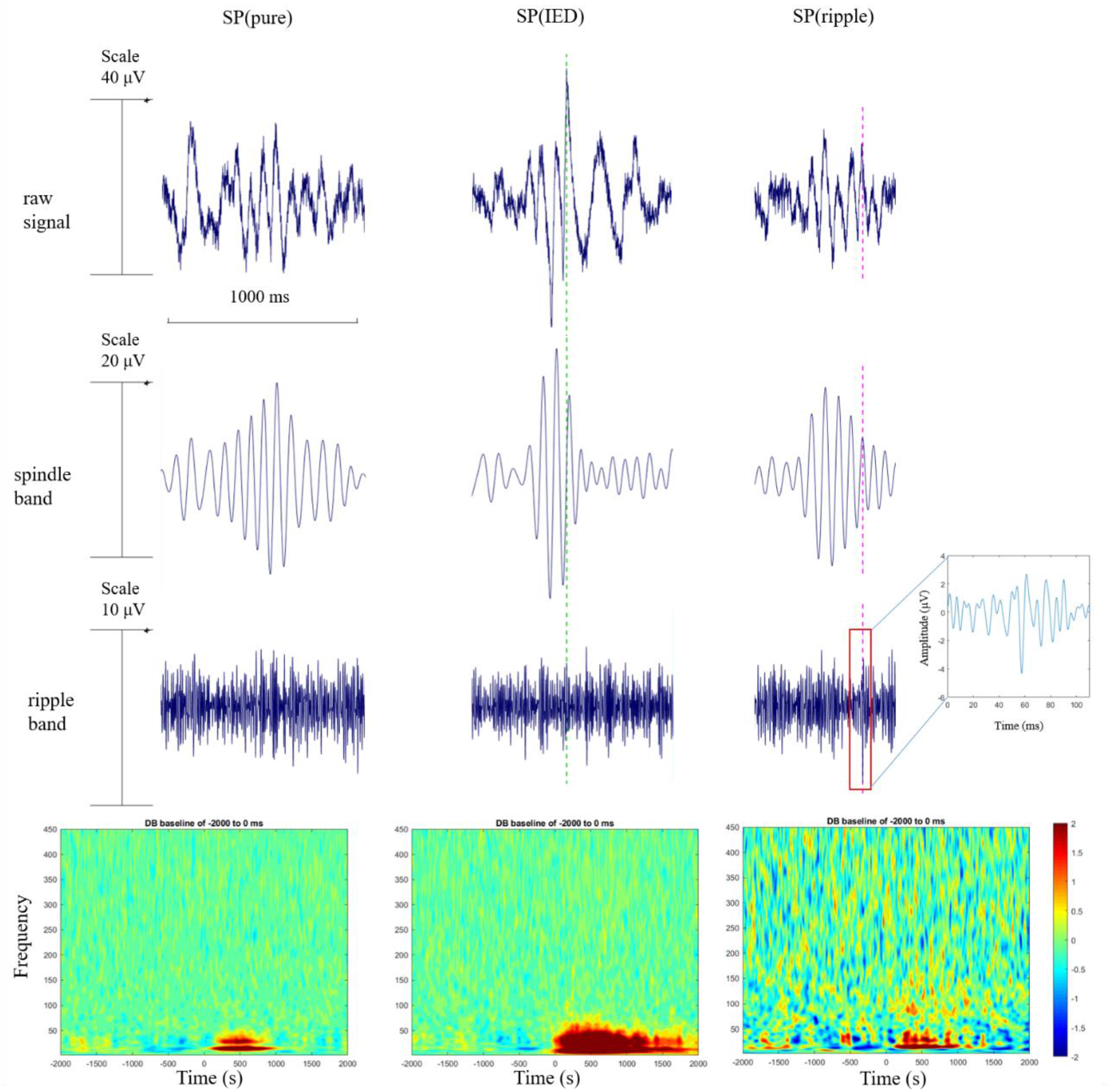
Examples of sleep spindles detected in the ANT from Patient #10: fast SP(pure) (left), SP(IED) (middle) and SP(ripple) (right). Upper row: 5 Hz highpass-filtered signal (raw signal), second row: 10–16 Hz bandpass-filtered signal (spindle band), third row: 80–200 Hz bandpass filtered signal (ripple band) of the same sleep spindle within each category. The fourth row depicts the time-frequency power spectra averaged across all fast sleep spindles of the same category detected in one ANT derivation of the same patient. The timepoint “0” indicates the onset of sleep spindles.

**Figure 4.**
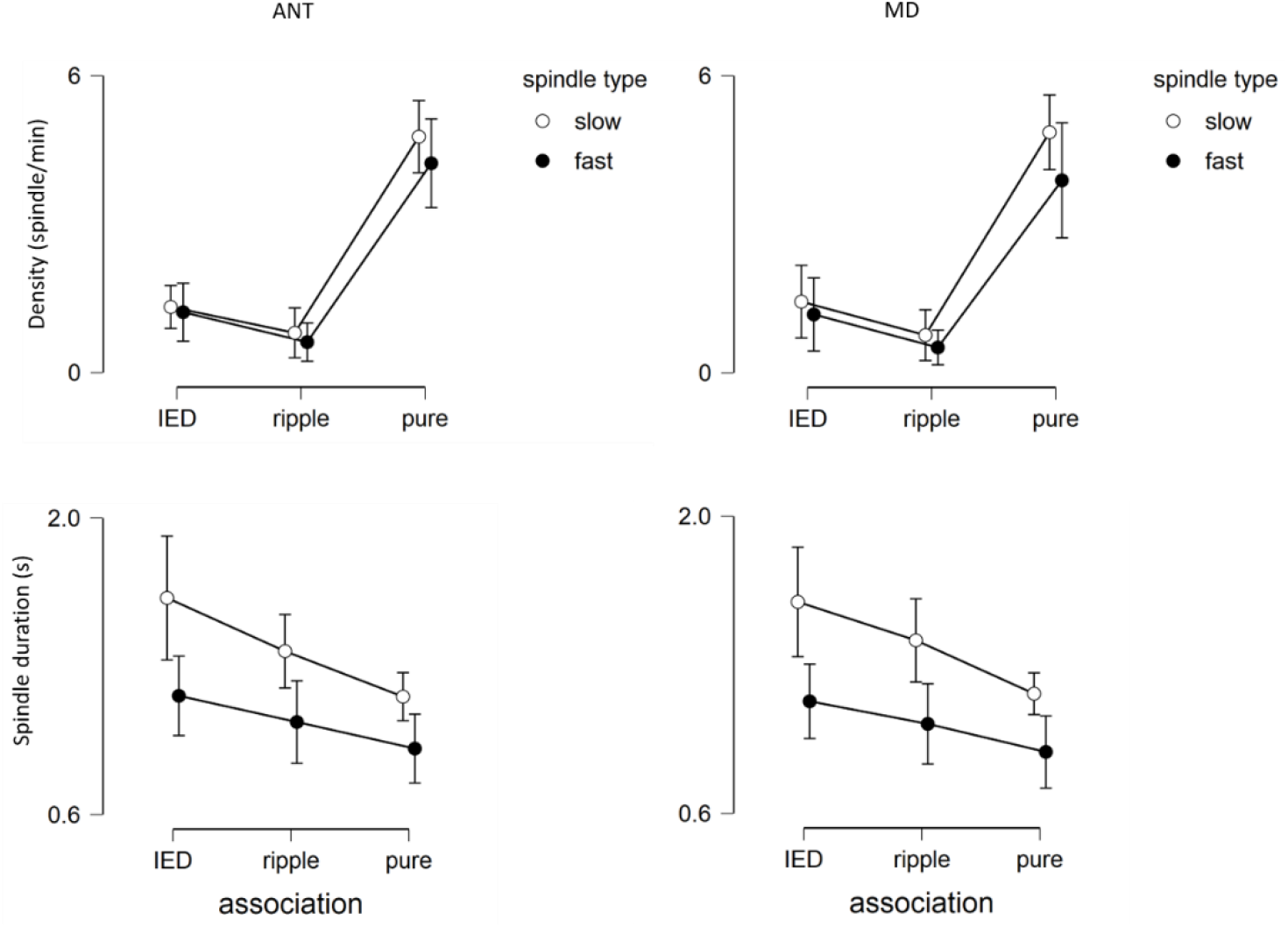
Sleep spindle density and duration in the ANT (left) and MD (right), separately for the IED associated (SP(IED)), ripple associated (SP(ripple)) and pure (SP(pure)) slow (white dots) and fast (black dots) spindles.

The duration of sleep spindles was affected by the ripple/IED association in both nuclei, as indicated by the significant main effects of association (F(2,28)=60.736, p<0.001, ε=0.697, η_p_^2^=0.813 for the ANT and F(2,18)= 38.118, p<0.001, ε=0.605, η_p_^2^= 0.809 for the MD): SP(ripple) and SP(IED) were longer than SP(pure), (p<0.001, all) and SP(IED) were also longer than SP(ripple), (ANT: p<0.001, MD: p=0.004). The main effect of spindle type was also significant, as longer slow spindles were detected compared to the fast spindles (F(1,14)=6.114, p=0.027, η_p_^2^=0.304 for the ANT and F(1,9)= 6.893, p<0.001, η_p_^2^= 0.434 for the MD). Furthermore, the interaction between spindle type and association was significant in the ANT (F(2,28)=60.736, p<0.001, ε=0.697, η_p_^2^=0.813), indicating that the association of ripples and IEDs with slow spindles resulted more elongated spindles than in the case of fast spindles.

#### Time-frequency power spectra analysis of pure, IED-associated and ripple-associated spindles

SP(pure) and SP(ripple) were characterized by comparable spindle frequency power (11–16 Hz) in both thalamic nuclei (ANT and MD), while the 80–200 Hz power of ANT slow and fast SP(ripple) exceeded the corresponding power values of SP(pure) up to around 1000 ms (Figure 7). In the MD, the same contrast resulted significant differences for slow spindles, but not for the fast spindles. Furthermore, slow SP(ripple) in the MD were associated with significantly higher theta power in the 980–1500 ms time window. Thalamic SP(pure) and SP(ripple) did not differ in terms of concomitant scalp EEG time-frequency power (see Supplementary figure 1 and 2). In contrast, significantly higher broad-band power ranging from the delta, up to the fast ripple frequencies were revealed during thalamic SP(IED) (time window: from spindle onset up to 1500 ms), in both nuclei (Figure 5).

**Figure 5.**
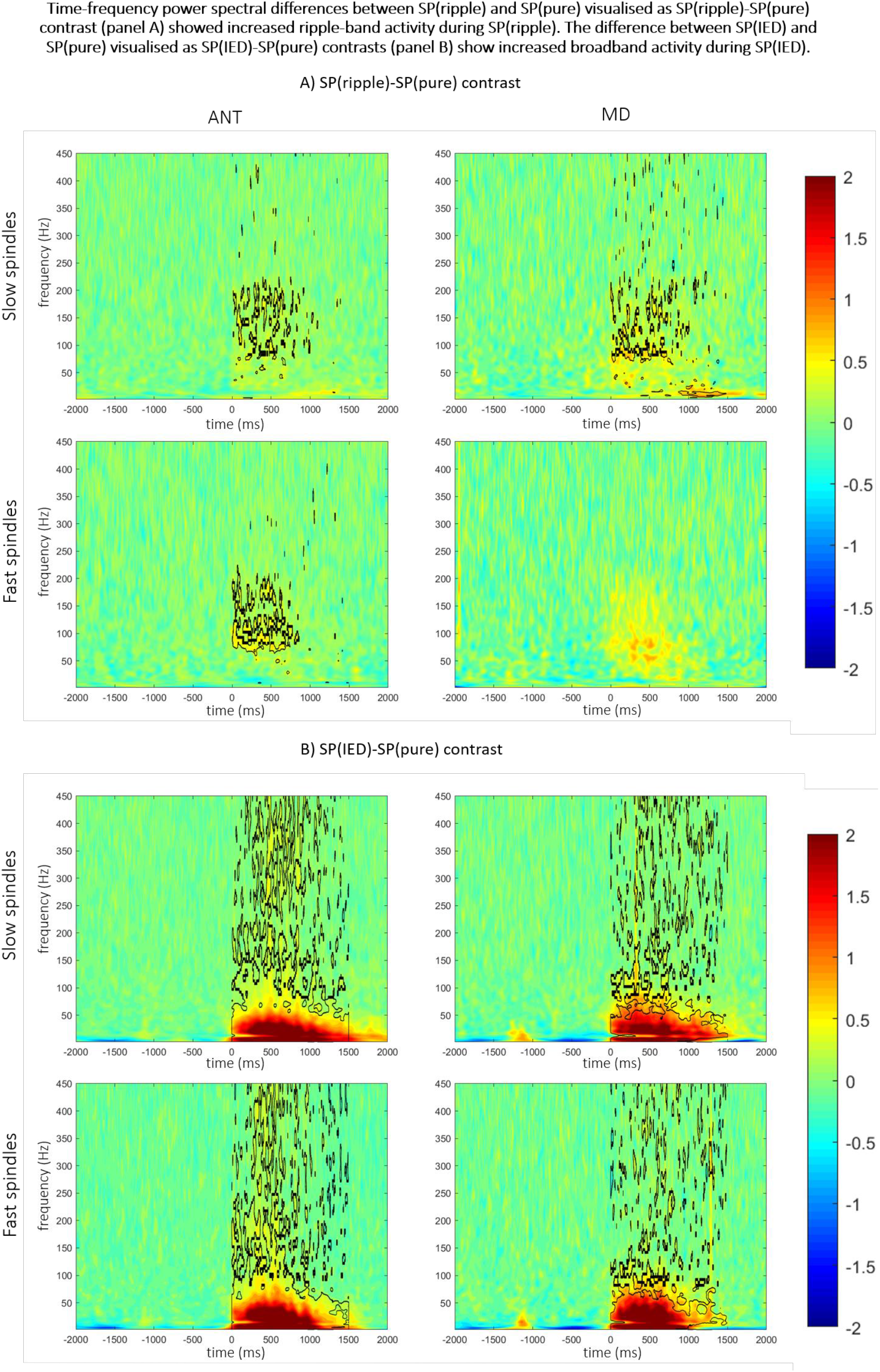
Time-frequency power spectra differences between SP(ripple) and SP(pure), and SP(IED) and SP(pure), separately for the ANT (left panel) and MD (right panel), and slow (upper row) and fast spindle (bottom row) within the 1–450 Hz frequency range. The timepoint “0” indicates the onset of sleep spindles. Significant time-frequency bins are circumscribed by the black lines.

Excess broadband ANT and MD LFP power during thalamic SP(IED) was associated with concomitant increases in scalp EEG delta, theta, beta and gamma, but not spindle and ripple activity (Figure 6).

**Figure 6.**
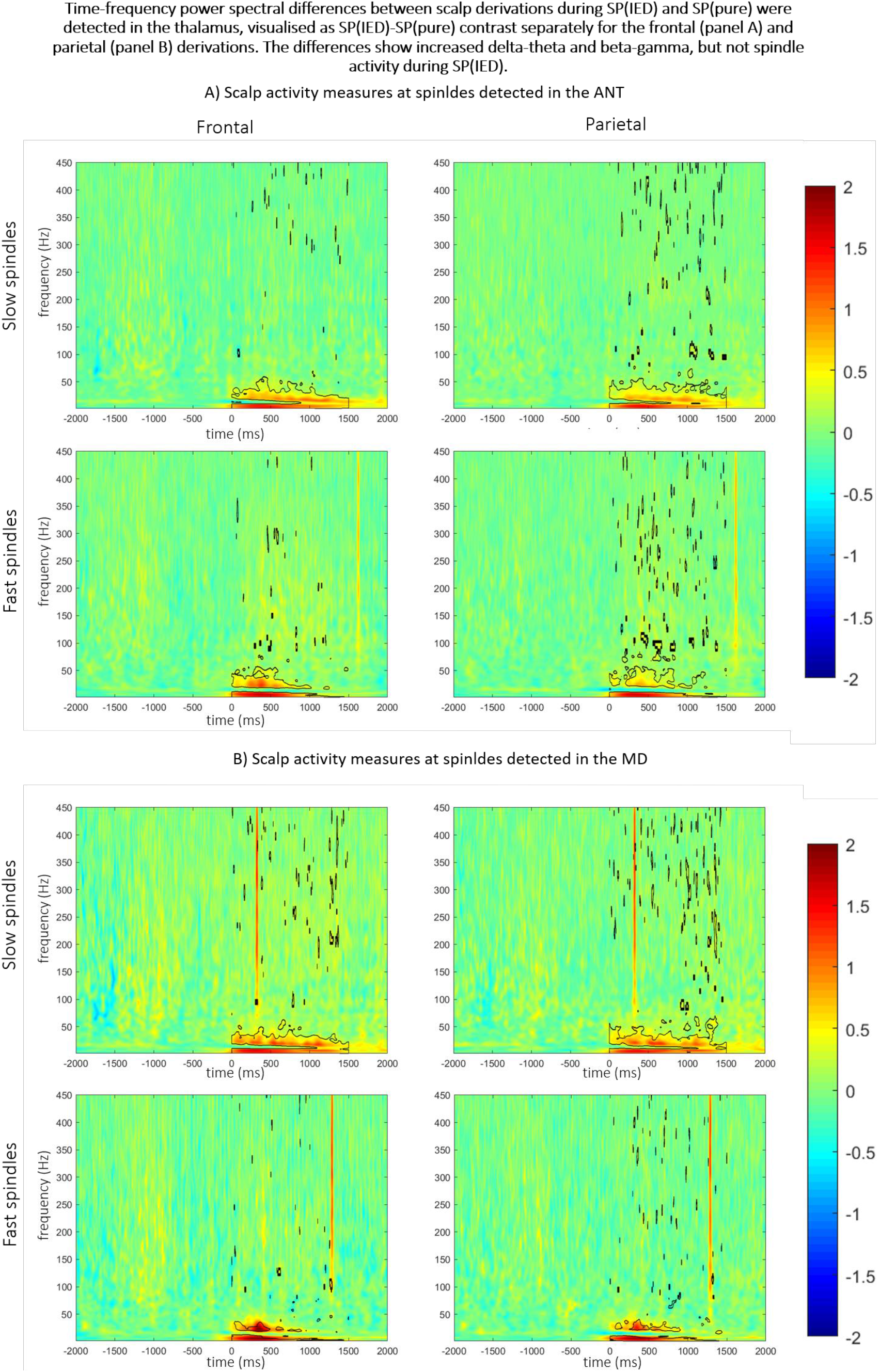
Time-frequency scalp EEG power spectra differences between scalp measures during ANT (panel A) and MD (panel B) SP(IED) and SP(pure) for frontal (left panel) and parietal (right panel) recordings, as well as slow (upper row) and fast spindle (bottom row) events within the 1–450 Hz frequency range. The timepoint “0” indicates the onset of ANT and MD sleep spindles. Significant time-frequency bins are circumscribed by the black lines.

#### Spindle dynamics in the thalamus and the cortex

Finally, we aimed to estimate the time dynamics of spindle co-occurrence between thalamic and cortical channels. Spindle co-occurrence was defined in the following way: when the initiation of a sleep spindle was detected on any channel (cortical or thalamic), all subsequent spindles initiating on any other channel before the end of the original spindle were considered to co-occur, comprising a single spindle event involving multiple channels with a time lag on each channel, defined as the time difference of spindle initiation relative to the first spindle.

For thalamocortical co-occurrence analysis, we selected all instances when sleep spindles co-occurred on both 1) a selected scalp channel (F3 or F4 for frontal spindles, P3 or P4 for parietal spindles, for patient #2 Fp2-Fpz and Pz-Oz instead) and 2) on a channel localized in a specific thalamic nucleus (ANT or MD). That is, the analysis included spindles which originated elsewhere but were later detected on both specific scalp channels and in the thalamus. We defined thalamocortical spindle lags as the time lag (relative to the first spindle) of the scalp channel minus the time lag of the thalamic channel.

The distribution of thalamocortical lags (broken down by ripple or IED occurrence) is shown on Figure 7. In short, thalamocortical lags were generally either negative or not significantly different from zero, indicating that either sleep spindles preferentially occurred on the scalp first or occurred simultaneously on scalp and thalamic channels.

**Figure 7.**
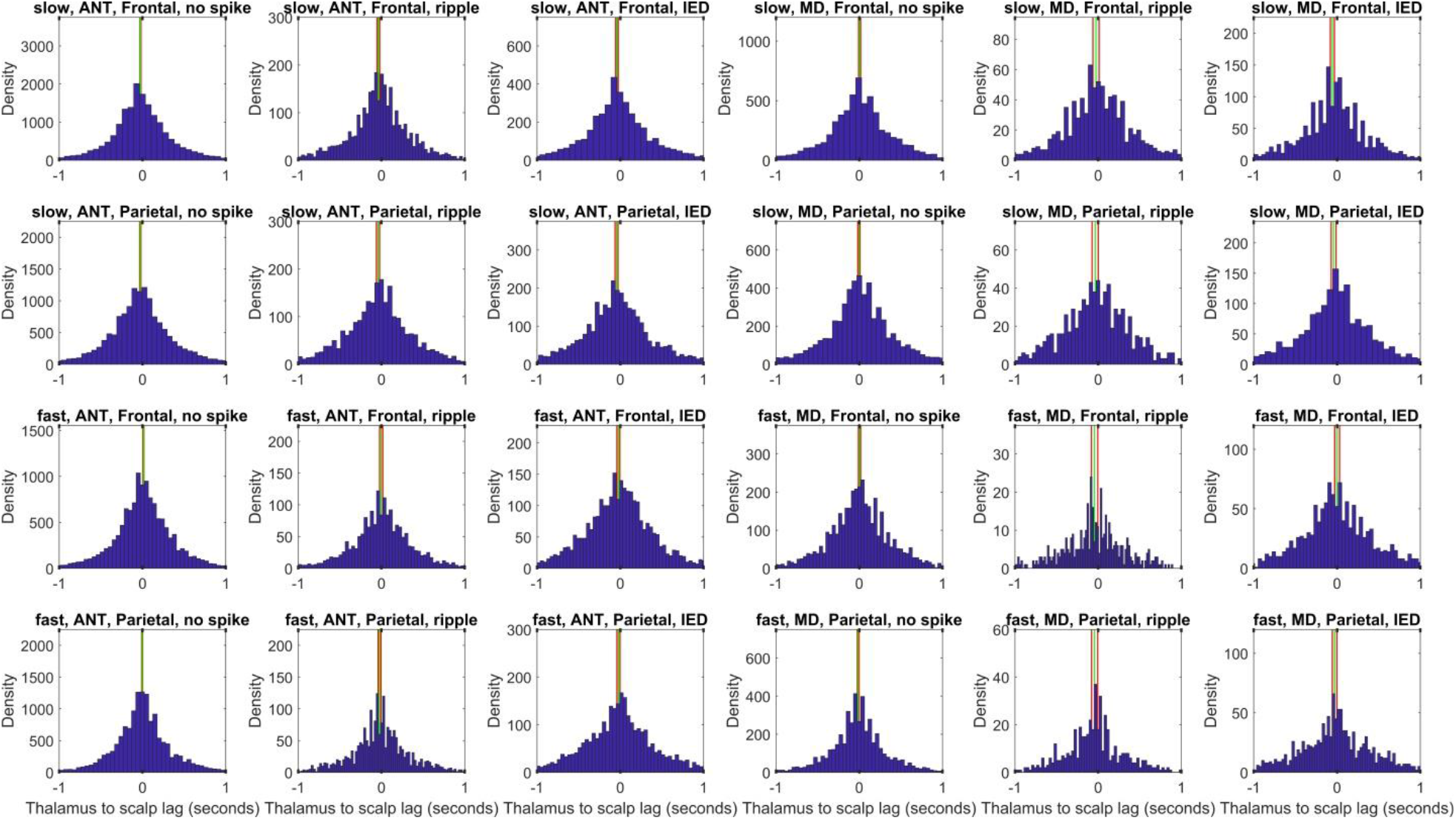
Thalamocortical spindle lags by ripple or IED occurrence. Histograms show the time lag difference of scalp minus thalamic spindles by scalp region, thalamic nucleus, IED presence and spindle type (slow or fast), pooled across all patients. Vertical green lines show the mean lag and red lines indicate the 95% CIs. Negative values indicate that spindles preferentially occur on the scalp first and vice versa. Axis X on each histogram is truncated at [-1 1] seconds for optimal visibility.

For a more precise analysis, we modeled spindle lags with a linear mixed model implemented in the MATLAB 2017a fitlme() function using lag as the dependent variable, spindle type, thalamic nucleus and scalp channel as fixed effects with random intercepts by patient. Very little variance in spindle lag was accounted for by this model (R2=0.002), suggesting that lag mainly depends on extraneous factors. Still, fast spindles (β=0.018), spindles in the MD nucleus (β=0.012) and spindles occurring on F3 (β=0.009) were associated with more positive lags, while spindles occurring on P3 (β=-0.007) and spindles with either ripples (β=-0.021) or IEDs (β=-0.024) were associated with more negative lags. That is, in case of fast spindles, in the MD and on F3, there was a diminished tendency for scalp spindles to occur first, but this tendency increased on P3 and especially when epileptiform activity or ripple activity were present in the thalamus. (All effect sizes are unstandardized and given in seconds. All p<0.001, except for the effect associated with F3 [p=0.005]).

### 3. The relation between sleep spindle types and general intelligence

The overall density of sleep spindles did not show significant correlation with IQ, neither in the ANT (r = 0.139, p = 0.682 for slow and r = 0.029, p = 0.932 for fast sleep spindles, N = 11), nor in the MD (r = 0.311, p = 0.497 for slow and r = -0.523, p = 0.229 for fast sleep spindles, N = 7). Furthermore, fast spindle density measured on the scalp did not reach the specified alpha level (r = 0.399, p = 0.225). However, when thalamic sleep spindles were separated according to their associations with IEDs and ripples, positive correlations were observed between IQ and slow and fast SP(ripple) density in the ANT (Figure 8; r = 0.661, N = 11, p=0.027 for the slow and r = 0.851, N = 11, p<0.001 for the fast spindles), but not in the MD (r = 0.703, N = 7, p=0.078 for the slow and r = 0.568, N = 7, p=0.184 for the fast spindles).

**Figure 8.**
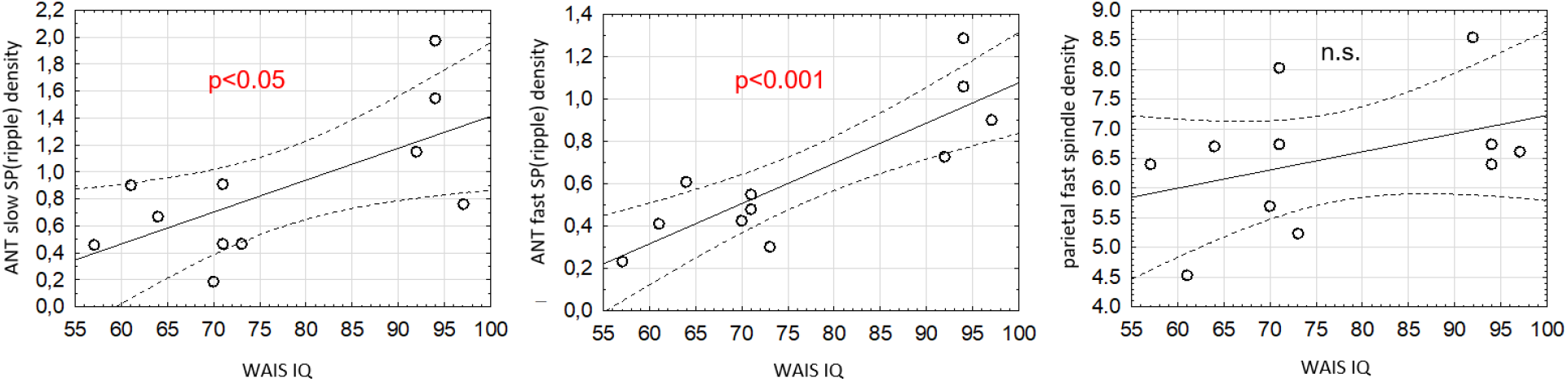
Correlations between general intelligence (IQ) and ripple associated slow (left) and fast (middle) sleep spindles in the ANT, and parietal scalp fast spindles (right).

## Discussion

We investigated the occurrence of sleep spindles, ripples and interictal epileptiform discharges in the human anterior and mediodorsal thalamus and those associations with epilepsy characteristics and general intelligence. Sleep spindles were detected both in the ANT and MD. The occurrence of sleep spindles in the human ANT were confirmed in previous studies (Tsai et al., 2010). We demonstrated that spindles are also present in the human MD. The overall density and duration of slow and fast spindles were similar in the two thalamic nuclei. In the majority of cases where thalamic-scalp sleep spindle co-occurrences were detected, ANT and MD sleep spindles slightly lagged behind cortical ones, which coheres with available reports (Tsai et al., 2010) and suggests the prevailing role of corticofugal fibers in the herein studied phenomena. In a recent study, Bastuji et al. (2020) recorded sleep spindles from different parts of the posterior thalamus.

These results indicate that anterior, mediodorsal and posterior thalamic nuclei are involved in the proper functioning of the NREM sleep-related thalamo-cortical network generating mid-frequency oscillations in the burst-firing mode.

In addition of detecting thalamic sleep spindles in human subjects, we revealed their association with thalamic IEDs and ripples. SP(IED) and SP(ripple) made up around 20 and 10% of overall ANT and MD sleep spindles, respectively. Both of these spindles were of longer duration although characterized by different time-frequency activity profiles. That is, thalamic IEDs preferentially emerged during ongoing sleep spindles, whereas thalamic ripple occurrence did not differentiate spindle and non-spindle NREM epochs. SP(IED) were characterized by an excess of slow and high frequency activity in both thalamic LFP and scalp EEG, whereas SP(ripple) were only and exclusively associated with an excess of thalamic ripple frequency activity (which was the basis of their definition). No cortical (scalp-recordable) signature of thalamic SP(ripple) were revealed. Furthermore, SP(IED) occurrence rate in the thalamic LFP correlated with visually-detected thalamic IED density, but no such association was evident for ripples. IED(sp), but not ripple(sp) density in the MD were shown to associate with years since epilepsy onset. Last, but not least ANT SP(ripple) were shown to indicate preserved cognitive ability. These findings suggest that thalamic ripples and IEDs could indicate physiological and pathological forms of neural plasticity/excitation, respectively. Moreover, our findings cohere with reports supporting the relevance of the thalamus in epilepsy.

Former studies suggested that IEDs in the human ANT contribute to the epileptic network, and the ANT became an important target for DBS in epilepsy treatment (Hodaie et al., 2002; Salanova, 2018; Sweeney-Reed et al., 2016). Furthermore, interictal discharges were also detected in the MD (Sweeney-Reed et al., 2016). Although, there is no direct anatomical connection between the ANT and MD, this report indicates that the MD might be involved in the epileptic circuitry as well. Our current results further support this assumption. Besides IEDs, the sleep spindle types (slow vs fast) could be also indicative for specific network dysfunctions. The density of slow SP(pure) in the MD showed a negative correlation with the year since epilepsy onset. In contrast to slow spindles, fast SP(pure) density showed a positive correlation with seizure prevalence. These findings suggest that the ANT and MD are involved in different epileptogenic circuitries. The occurrence of fast sleep spindles may indicate the altered functioning of the MD due to network-based changes through epilepsy propagation, whereas the occurrence of fast spindles seems physiological in the ANT. Sleep spindles in the mediodorsal thalamus were also indicative in schizophrenia patients, where a negative association were found between the volume of the MD and scalp-recorded sleep spindle density (for a review, see Ferrarelli & Tononi, 2017). To our best knowledge, this is the first report where sleep spindles were separated to fast and slow spindles, showing that the appearance of fast sleep spindles may indicate pathological mechanisms in the mediodorsal thalamic functions.

In the prefrontal cortex, mesio-temporal and neocortical structures, ripples were detected both before sleep spindles and locked to the spindle troughs (Bruder et al., 2021; Peyrache et al., 2011). In the current study, ripples were found in both thalamic nuclei coupled with sleep spindles, and outside spindles during NREM sleep. The coordinated interactions between hippocampal ripples and cortical sleep spindles plays crucial role in memory formation (Latchoumane et al., 2017; Siapas & Wilson, 1998; Staresina et al., 2015). It was proposed that NREM sleep-based memory formation depends on the hierarchical nesting of slow waves, sleep spindles and hippocampal ripples, as these oscillations are instrumental in transferring information from the hippocampus to the neocortex (Diekelmann & Born, 2010; Staresina et al., 2015). Although it is not entirely clear whether high frequency oscillations (HFO) in the thalamus are connected with the pathomechanism of epilepsy, a close relationship between epilepsy and HFO (80–600 Hz) occurrence is repeatedly found in animal and human studies (for a review, see Jiruska et al., 2017). In a recent study, Rektor et al. (2016) found HFOs in the human ANT up to 240 Hz frequency, and 500 Hz frequency in one case, which was the first report of HFOs in the human thalamus. They suspected that thalamic HFOs are related to pathological processes. Here we found a positive correlation between SP(ripple) and general intelligence in the ANT, which may indicate that ripples at lower frequencies (80–200 Hz) in the human ANT may contribute to physiological ripples which play important role in IQ and perhaps memory functioning. Sleep spindles, and especially fast sleep spindles measured on the scalp have a close connection with cognitive performance and intelligence (Bestmann et al., 2019; Bódizs et al., 2005; Chatburn et al., 2013; Ujma, 2018b). The ANT has tight anatomical and functional connection with the hippocampus, as well as different parts of the prefrontal cortex, and might act as an interface between them during memory consolidation. Although there is no direct measurement of memory performance in the current study, the results suggest that higher intelligence score is also associated with preserved memory performance, in which anterior thalamic ripple-spindle coupling might be involved.

It was found that cortical sleep spindles show a tendency of elongation when IEDs were detected in the hippocampus ca. 1 s prior to sleep spindles (Gelinas et al., 2016). Furthermore, the spectro-temporal characteristics of cortical sleep spindles preceded with IEDs were the same as those of physiological spindles. In contrast, we found that when IEDs are coupled with spindles in the thalamus, both slow and fast spindles became significantly longer than physiological sleep spindles: Sleep spindles coupled with IEDs (SP(IED)) were longer compared to those associated with ripples (SP(ripple)) and pure (SP(pure)) spindles in both nuclei. Furthermore, the time-frequency spectro-temporal characteristics also showed clear difference in the spectral power of SP(IED) compared to SP(pure), not only in the spindle frequency band but also at lower and higher frequencies, up to the frequency of HFOs. Thus, SP(IED) can be distinguished from physiological spindles based on a clear spectro-temporal difference between them. Furthermore, this spectro-temporal difference is also measurable at the scalp, at the same time interval when the sleep spindles are detected in the thalamus. This scalp difference occurs predominantly at lower frequencies (delta-theta) and in the beta frequency band, but only sporadically present at the spindle and high (>40 Hz) frequency band. Furthermore, the temporal dynamics of the spindles in the thalamus and the scalp suggest that cortical spindle preceded thalamic spindles associated with IEDs, and also with ripples. It was shown that IEDs induce strong and synchronized neural firing in the mPFC associated with the expression of delta waves and spindles, even outside NREM sleep (Kleen et al., 2011). The thalamus seems to play an important interface between the propagation of these oscillations in the hippocampal-prefrontal network. In contrast to SP(IED), ripple coupling (SP(ripple)) do not result in significant spectro-temporal differences in the scalp-measured oscillations, and reasonable difference in the thalamus was found only in the ripple (80–200 Hz) band. Although, the occurrence of ripples also elongated the sleep spindles, this result does not indicate any pathological mechanisms, instead, as reported above, spindles coupled with ripples might index intellectual ability.

The current results support the involvement of the human thalamus in sleep spindle-related neural plasticity. Sleep spindles were found in the ANT and in the MD and IEDs coupled with sleep spindles resulted in distinguishable spectro-temporal differences in the thalamus and also at the scalp. The major finding is that high frequency oscillations (ripples) are also present in the thalamus, which seems to contribute to intellectual ability and memory performance through a tight interaction between spindles and ripples. Furthermore, interictal discharges interfere with this process resulting pathological expression of sleep spindles. Limitations of the study is no direct testing of offline memory processes during sleep. Co-registration from the hippocampus, thalamus and scalp will present an opportunity to reveal the hippocampal-thalamo-cortical pathway in further research, providing target circuits for neuromodulation and therapeutics.

## References

Aggleton, J. P., O’Mara, S. M., Vann, S. D., Wright, N. F., Tsanov, M., & Erichsen, J. T. (2010). Hippocampal-anterior thalamic pathways for memory: Uncovering a network of direct and indirect actions. European Journal of Neuroscience, 31(12), 2292–2307. https://doi.org/10.1111/j.1460-9568.2010.07251.x

Bastuji, H., Lamouroux, P., Villalba, M., Magnin, M., & Garcia-Larrea, L. (2020). Local sleep spindles in the human thalamus. The Journal of Physiology, 598(11), 2109–2124. https://doi.org/10.1113/JP279045

Berry, R. B., Brooks, R., Gamaldo, C. E., Harding, S. M., Lloyd, R. M., Marcus, C. L., & Vaughn, B. V. (2015). AASM | Scoring Manual Version 2.2 The AASM Manual for the Scoring of Sleep and Associated Events Rules, Terminology And Technical Specifications Version 2.2. www.aasmnet.org.

Bestmann, A., Conzelmann, A., Baving, L., & Prehn-Kristensen, A. (2019). Associations between cognitive performance and sigma power during sleep in children with attention-deficit/hyperactivity disorder, healthy children, and healthy adults. PLOS ONE, 14(10), e0224166. https://doi.org/10.1371/journal.pone.0224166

Bódizs, R., Kis, T., Lázár, A. S., Havrán, L., Rigó, P., Clemens, Z., & Halász, P. (2005). Prediction of general mental ability based on neural oscillation measures of sleep. Journal of Sleep Research, 14(3), 285–292. https://doi.org/10.1111/j.1365-2869.2005.00472.x

Bódizs, R., Körmendi, J., Rigó, P., & Lázár, A. S. (2009). The individual adjustment method of sleep spindle analysis: Methodological improvements and roots in the fingerprint paradigm. Journal of Neuroscience Methods, 178(1), 205–213. https://doi.org/10.1016/j.jneumeth.2008.11.006

Bruder, J. C., Schmelzeisen, C., Lachner-Piza, D., Reinacher, P., Schulze-Bonhage, A., & Jacobs, J. (2021). Physiological Ripples Associated With Sleep Spindles Can Be Identified in Patients With Refractory Epilepsy Beyond Mesio-Temporal Structures. Frontiers in Neurology, 0, 48. https://doi.org/10.3389/FNEUR.2021.612293

Buchmann, A., Dentico, D., Peterson, M. J., Riedner, B. A., Sarasso, S., Massimini, M., Tononi, G., & Ferrarelli, F. (2014). Reduced mediodorsal thalamic volume and prefrontal cortical spindle activity in schizophrenia. NeuroImage, 102(P2), 540–547. https://doi.org/10.1016/j.neuroimage.2014.08.017

Burgaleta, M., MacDonald, P. A., Martínez, K., Román, F. J., Álvarez-Linera, J., González, A. R., Karama, S., & Colom, R. (2014). Subcortical regional morphology correlates with fluid and spatial intelligence. Human Brain Mapping, 35(5), 1957–1968. https://doi.org/10.1002/hbm.22305

Cairney, S. A., Guttesen, A. V., Marj, N. E., & Staresina, B. P. (2018). Memory Consolidation Is Linked to Spindle-Mediated Information Processing during Sleep. Current Biology, 28(6), 948. https://doi.org/10.1016/J.CUB.2018.01.087

Chatburn, A., Coussens, S., Lushington, K., Kennedy, D., Baumert, M., & Kohler, M. (2013). Sleep Spindle Activity and Cognitive Performance in Healthy Children. Sleep, 36(2), 237–243. https://doi.org/10.5665/sleep.2380

Child, N. D., & Benarroch, E. E. (2013). Anterior nucleus of the thalamus:Functional organization and clinical implications. Neurology, 81(21), 1869–1876. https://doi.org/10.1212/01.wnl.0000436078.95856.56

Deutschová, B., Klimeš, P., Jordan, Z., Jurák, P., Erőss, L., Lamoš, M., Halámek, J., Daniel, P., Rektor, I., & Fabo, D. (2021). Thalamic oscillatory activity may predict response to deep brain stimulation of the anterior nuclei of the thalamus. Epilepsia, 62(5), e70–e75. https://doi.org/10.1111/epi.16883

Diekelmann, S., & Born, J. (2010). The memory function of sleep. Nature Reviews Neuroscience, 11(2), 114–126. https://doi.org/10.1038/nrn2762

Ferrarelli, F., & Tononi, G. (2017). Reduced sleep spindle activity point to a TRN-MD thalamus-PFC circuit dysfunction in schizophrenia. Schizophrenia Research, 180, 36–43. https://doi.org/10.1016/j.schres.2016.05.023

Fogel, S. M., & Smith, C. T. (2011). The function of the sleep spindle: A physiological index of intelligence and a mechanism for sleep-dependent memory consolidation. Neuroscience and Biobehavioral Reviews, 35(5), 1154–1165. https://doi.org/10.1016/j.neubiorev.2010.12.003

Gelinas, J. N., Khodagholy, D., Thesen, T., Devinsky, O., & Buzsáki, G. (2016). Interictal epileptiform discharges induce hippocampal-cortical coupling in temporal lobe epilepsy. Nature Medicine, 22(6), 641–648. https://doi.org/10.1038/nm.4084

Girardeau, G., & Zugaro, M. (2011). Hippocampal ripples and memory consolidation. Current Opinion in Neurobiology, 21(3), 452–459. https://doi.org/10.1016/j.conb.2011.02.005

Halász, P., Bódizs, R., Ujma, P. P., Fabó, D., & Szűcs, A. (2019). Strong relationship between NREM sleep, epilepsy and plastic functions — A conceptual review on the neurophysiology background. Epilepsy Research, 150, 95–105. https://doi.org/10.1016/j.eplepsyres.2018.11.008

Halász, P., & Szűcs, A. (2020). Sleep and Epilepsy Link by Plasticity. Frontiers in Neurology, 11, 911. https://doi.org/10.3389/FNEUR.2020.00911

Hodaie, M., Wennberg, R. A., Dostrovsky, J. O., & Lozano, A. M. (2002). Chronic Anterior Thalamus Stimulation for Intractable Epilepsy. Epilepsia, 43(6), 603–608. https://doi.org/10.1046/j.1528-1157.2002.26001.x

Huguenard, J. R., & McCormick, D. A. (2007). Thalamic synchrony and dynamic regulation of global forebrain oscillations. Trends in Neurosciences, 30(7), 350–356. https://doi.org/10.1016/j.tins.2007.05.007

Jasper, H. H. (1958). The Ten-Twenty Electrode System of the International Federation. Electroencephalography and Clinical Neurophysiology, 10, 371–375.

Jiruska, P., Alvarado-Rojas, C., Schevon, C. A., Staba, R., Stacey, W., Wendling, F., & Avoli, M. (2017). Update on the mechanisms and roles of high-frequency oscillations in seizures and epileptic disorders. Epilepsia, 58(8), 1330–1339. https://doi.org/10.1111/epi.13830

Kleen, J. K., Wu, E. X., Holmes, G. L., Scott, R. C., & Lenck-Santini, P. P. (2011). Enhanced oscillatory activity in the hippocampal-prefrontal network is related to short-term memory function after earlylife seizures. Journal of Neuroscience, 31(43), 15397–15406. https://doi.org/10.1523/JNEUROSCI.2196-11.2011

Latchoumane, C. F. V., Ngo, H. V. V., Born, J., & Shin, H. S. (2017). Thalamic spindles promote memory formation during sleep through triple phase-locking of cortical, thalamic, and hippocampal rhythms. Neuron, 95(2), 424-435.e6. https://doi.org/10.1016/j.neuron.2017.06.025

Lugaresi, E., Tobler, I., Gambetti, P., & Montagna, P. (2006). The pathophysiology of fatal familial insomnia. Brain Pathology, 8(3), 521–526. https://doi.org/10.1111/j.1750-3639.1998.tb00173.x

Mak-Mccully, R. A., Rolland, M., Sargsyan, A., Gonzalez, C., Magnin, M., Chauvel, P., Rey, M., Bastuji, H., & Halgren, E. (2017). Coordination of cortical and thalamic activity during non-REM sleep in humans. Nature Communications, 8(1), 1–11. https://doi.org/10.1038/ncomms15499

Manzano, G. M., Ragazzo, P. C., Tavares, S. M., & Marino Jr., R. (1986). Anterior Zygomatic Electrodes: A Special Electrode for the Study of Temporal Lobe Epilepsy. Stereotactic and Functional Neurosurgery, 49(4), 213–217. https://doi.org/10.1159/000100148

Mitchell, A. S., & Chakraborty, S. (2013). What does the mediodorsal thalamus do? Frontiers in Systems Neuroscience, JUL. https://doi.org/10.3389/fnsys.2013.00037

Oostenveld, R., Fries, P., Maris, E., & Schoffelen, J. M. (2011). FieldTrip: Open source software for advanced analysis of MEG, EEG, and invasive electrophysiological data. Computational Intelligence and Neuroscience, 2011. https://doi.org/10.1155/2011/156869

Peyrache, A., Battaglia, F. P., & Destexhe, A. (2011). Inhibition recruitment in prefrontal cortex during sleep spindles and gating of hippocampal inputs. Proceedings of the National Academy of Sciences of the United States of America, 108(41), 17207–17212. https://doi.org/10.1073/pnas.1103612108

Rektor, I., Doležalová, I., Chrastina, J., Jurák, P., Halámek, J., Baláž, M., & Brázdil, M. (2016). High-Frequency Oscillations in the Human Anterior Nucleus of the Thalamus. Brain Stimulation, 9(4), 629–631. https://doi.org/10.1016/j.brs.2016.04.010

Salanova, V. (2018). Deep brain stimulation for epilepsy. Epilepsy and Behavior, 88, 21–24. https://doi.org/10.1016/j.yebeh.2018.06.041

Saletin, J. M., Goldstein, A. N., & Walker, M. P. (2011). The Role of Sleep in Directed Forgetting and Remembering of Human Memories. Cerebral Cortex (New York, NY), 21(11), 2534. https://doi.org/10.1093/CERCOR/BHR034

Siapas, A. G., & Wilson, M. A. (1998). Coordinated interactions between hippocampal ripples and cortical spindles during slow-wave sleep. Neuron, 21(5), 1123–1128. https://doi.org/10.1016/S0896-6273(00)80629-7

Simor, P., Szalárdy, O., Gombos, F., Ujma, P. P., Jordán, Z., Halász, L., Erőss, L., Fabó, D., & Bódizs, R. (2021). REM Sleep Microstates in the Human Anterior Thalamus. The Journal of Neuroscience, JN-RM-1899-20. https://doi.org/10.1523/JNEUROSCI.1899-20.2021

Staresina, B. P., Bergmann, T. O., Bonnefond, M., Van Der Meij, R., Jensen, O., Deuker, L., Elger, C. E., Axmacher, N., & Fell, J. (2015). Hierarchical nesting of slow oscillations, spindles and ripples in the human hippocampus during sleep. Nature Neuroscience, 18(11), 1679–1686. https://doi.org/10.1038/nn.4119

Steriade, M. (2005). Sleep, epilepsy and thalamic reticular inhibitory neurons. Trends in Neurosciences, 28(6 SPEC. ISS.), 317–324. https://doi.org/10.1016/j.tins.2005.03.007

Sweeney-Reed, C. M., Lee, H., Rampp, S., Zaehle, T., Buentjen, L., Voges, J., Holtkamp, M., Hinrichs, H., Heinze, H. J., & Schmitt, F. C. (2016). Thalamic interictal epileptiform discharges in deep brain stimulated epilepsy patients. Journal of Neurology, 263(10), 2120–2126. https://doi.org/10.1007/s00415-016-8246-5

Tsai, Y. T., Chan, H. L., Lee, S. T., Tu, P. H., Chang, B. L., & Wu, T. (2010). Significant thalamocortical coherence of sleep spindle, theta, delta, and slow oscillations in NREM sleep: Recordings from the human thalamus. Neuroscience Letters, 485(3), 173–177. https://doi.org/10.1016/j.neulet.2010.09.004

Ujma, P. P. (2018a). Sleep spindles and general cognitive ability – A meta-analysis. Sleep Spindles & Cortical Up States, 1, 1–17. https://doi.org/10.1556/2053.2.2018.01

Ujma, P. P. (2018b). Sleep spindles and general cognitive ability – A meta-analysis. Sleep Spindles & Cortical Up States, 1(aop), 1–17. https://doi.org/10.1556/2053.2.2018.01

Ujma, P. P., Bódizs, R., & Dresler, M. (2020). Sleep and intelligence: critical review and future directions. Current Opinion in Behavioral Sciences, 33, 109–117. https://doi.org/10.1016/j.cobeha.2020.01.009

Ujma, P. P., Gombos, F., Genzel, L., Konrad, B. N., Simor, P., Steiger, A., Dresler, M., & BÃ^3^dizs, R. (2015). A comparison of two sleep spindle detection methods based on all night averages: individually adjusted vs. fixed frequencies. Frontiers in Human Neuroscience, 9(FEB), 52. https://doi.org/10.3389/fnhum.2015.00052

Ujma, P. P., Simor, P., Ferri, R., Fabó, D., Kelemen, A., Eross, L., Bódizs, R., & Halász, P. (2015). Increased interictal spike activity associated with transient slow wave trains during non-rapid eye movement sleep. Sleep and Biological Rhythms, 13(2), 155–162. https://doi.org/10.1111/sbr.12101

Van Der Werf, Y. D., Jolles, J., Witter, M. P., & Uylings, H. B. M. (2003). Contributions of thalamic nuclei to declarative memory functioning. Cortex, 39(4–5), 1047–1062. https://doi.org/10.1016/S0010-9452(08)70877-3

