## Supplementary Material for "Sleep spindles, ripples, and interictal epileptiform discharges in the human anterior and mediodorsal thalamus"

The comparison of automatic IED and ripple detections revealed no significant correlations between IED(NREM) or IED(sp) density and manually detected mean thalamic or scalp IED density (see Supplementary Table 1.). However, slow and fast SP(IED) density in the ANT correlated positively with manually detected mean thalamic IED density, but not with mean scalp IED density. Furthermore, statistical analysis revealed a significant positive correlation between IED(sp) density in the MD and year since epilepsy onset. In contrast, slow SP(pure) density in the MD showed a negative correlation with the years since epilepsy onset, whereas the fast SP(pure) density in the same nucleus correlated positively with the seizure prevalence. No significant correlation occurred between any of the variables and age. Furthermore, no significant correlations were found between ripple(NREM and sp) density, and epilepsy characteristics, and between the SP(ripple) density and epilepsy characteristics. These results indicate that thalamic sleep spindles associated with IEDs (SP(IED)) might be reliable candidate biomarkers of pathologic, epilepsy-related plasticity. Thus, further statistical comparisons were performed on sleep spindle densities and durations, regarding the association with IEDs and ripples (SP(IED/ripple/pure)).

*Supplementary Table 1. Correlations between age, epilepsy, and manually detected IED density*

Correlations, marked correlations are significant at $p < .05$

| Variable | Age | years since epilepsy onset | seizure prevalence | Mean thalamic IED density (manually detected) | Mean scalp IED density (manually detected) |
|---|---|---|---|---|---|
| IED(sp) density ANT | 0.461<br>N=15<br>p=.084 | 0.466<br>N=15<br>p=.080 | -0.274<br>N=15<br>p=.324 | 0.341<br>N=15<br>p=.214 | 0.098<br>N=15<br>p=.729 |
| IED(NREM) density ANT | 0.209<br>N=15<br>p=.454 | -0.061<br>N=15<br>p=.829 | -0.233<br>N=15<br>p=.403 | -0.025<br>N=15<br>p=.931 | -0.050<br>N=15<br>p=.859 |
| IED(sp) density MD | 0.343<br>N=10<br>p=.332 | 0.714<br>N=10<br>p=.020 | -0.543<br>N=10<br>p=.105 | 0.262<br>N=10<br>p=.464 | -0.169<br>N=10<br>p=.642 |
| IED(NREM) density MD | 0.233<br>N=10<br>p=.518 | 0.440<br>N=10<br>p=.203 | -0.463<br>N=10<br>p=.178 | 0.229<br>N=10<br>p=.524 | -0.126<br>N=10<br>p=.730 |
| ripple(sp) density ANT | 0.417<br>N=15<br>p=.122 | 0.466<br>N=15<br>p=.080 | -0.273<br>N=15<br>p=.326 | 0.429<br>N=15<br>p=.111 | 0.088<br>N=15<br>p=.754 |
| ripple(NREM) density ANT | 0.095<br>N=15<br>p=.736 | -0.084<br>N=15<br>p=.766 | -0.162<br>N=15<br>p=.564 | -0.472<br>N=15<br>p=.076 | -0.436<br>N=15<br>p=.104 |
| ripple(sp) density MD | 0.288<br>N=10 | 0.613<br>N=10 | -0.486<br>N=10 | 0.323<br>N=10 | -0.162<br>N=10 |

| | | | | | |
|---|---|---|---|---|---|
| | p=.420 | p=.059 | p=.155 | p=.362 | p=.654 |
| ripple(NREM) density MD | 0.198 | 0.255 | -0.601 | -0.140 | -0.426 |
| | N=10 | N=10 | N=10 | N=10 | N=10 |
| | p=.583 | p=.478 | p=.066 | p=.701 | p=.220 |
| SP(IED) slow density ANT | 0.440 | 0.361 | -0.065 | 0.763 | 0.443 |
| | N=15 | N=15 | N=15 | N=15 | N=15 |
| | p=.101 | p=.186 | p=.819 | p=.001 | p=.098 |
| SP(IED) fast density ANT | 0.433 | 0.506 | -0.140 | 0.737 | 0.301 |
| | N=15 | N=15 | N=15 | N=15 | N=15 |
| | p=.107 | p=.054 | p=.619 | p=.002 | p=.276 |
| SP(IED) slow density MD | 0.340 | 0.493 | -0.433 | 0.343 | -0.025 |
| | N=10 | N=10 | N=10 | N=10 | N=10 |
| | p=.336 | p=.148 | p=.211 | p=.332 | p=.946 |
| SP(IED) fast density MD | 0.333 | 0.616 | -0.380 | 0.512 | 0.002 |
| | N=10 | N=10 | N=10 | N=10 | N=10 |
| | p=.347 | p=.058 | p=.278 | p=.131 | p=.995 |
| SP(ripple) slow density ANT | 0.024 | -0.104 | 0.042 | -0.169 | -0.164 |
| | N=15 | N=15 | N=15 | N=15 | N=15 |
| | p=.932 | p=.712 | p=.883 | p=.548 | p=.559 |
| SP(ripple) fast density ANT | 0.216 | 0.118 | -0.097 | -0.158 | -0.266 |
| | N=15 | N=15 | N=15 | N=15 | N=15 |
| | p=.441 | p=.676 | p=.731 | p=.573 | p=.338 |
| SP(ripple) slow density MD | 0.021 | -0.274 | -0.227 | -0.188 | -0.242 |
| | N=10 | N=10 | N=10 | N=10 | N=10 |
| | p=.954 | p=.444 | p=.529 | p=.604 | p=.500 |
| SP(ripple) fast density MD | 0.076 | -0.055 | -0.250 | -0.450 | -0.496 |
| | N=10 | N=10 | N=10 | N=10 | N=10 |
| | p=.835 | p=.880 | p=.486 | p=.192 | p=.145 |
| SP(pure) slow density ANT | -0.425 | -0.320 | 0.216 | 0.083 | 0.100 |
| | N=15 | N=15 | N=15 | N=15 | N=15 |
| | p=.114 | p=.245 | p=.440 | p=.768 | p=.723 |
| SP(pure) fast density ANT | -0.264 | -0.165 | 0.285 | 0.199 | 0.149 |
| | N=15 | N=15 | N=15 | N=15 | N=15 |
| | p=.342 | p=.558 | p=.304 | p=.477 | p=.597 |
| SP(pure) slow density MD | -0.304 | -0.731 | 0.617 | 0.364 | 0.345 |
| | N=10 | N=10 | N=10 | N=10 | N=10 |
| | p=.393 | p=.016 | p=.057 | p=.301 | p=.329 |
| SP(pure) fast density MD | -0.369 | -0.419 | 0.666 | -0.158 | 0.031 |
| | N=10 | N=10 | N=10 | N=10 | N=10 |
| | p=.294 | p=.228 | p=.035 | p=.662 | p=.932 |
| Mean thalamic IED density (manually detected) | 0.165 | | | | |
| | N=15 | | | | |
| | p=.557 | | | | |
| WAIS IQ | 0.236 | | | | |
| | N=11 | | | | |
| | p=.485 | | | | |

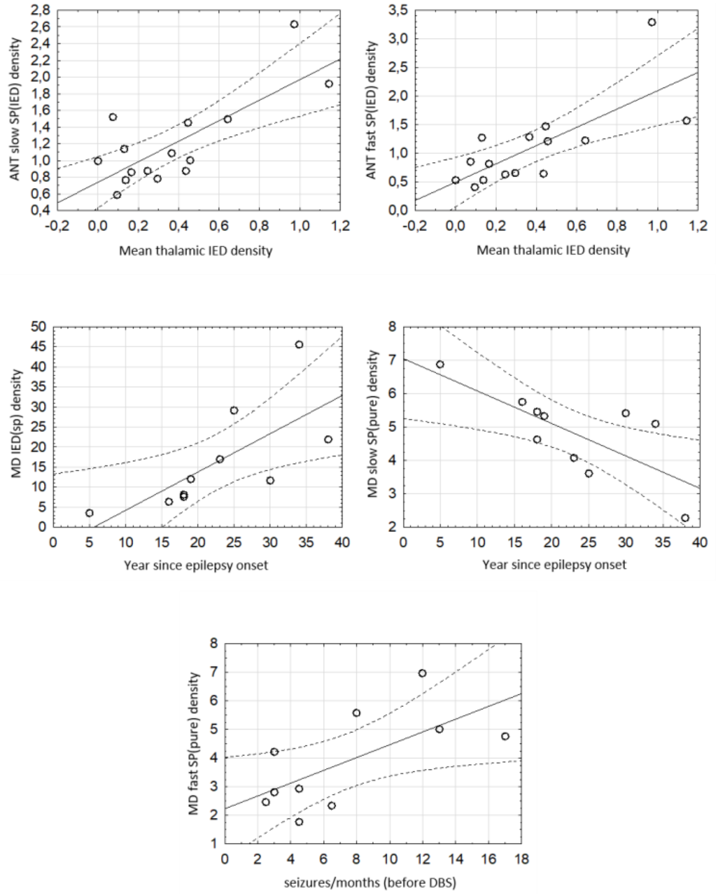


*Supplementary Figure 1. Correlations between epilepsy characteristics, IED density, and spindle densities. Top right and left panels show the significant positive correlations between ANT slow and fast SP(IED) densities and manually detected mean thalamic IED densities. Middle panels show the correlation between epilepsy duration (year since epilepsy onset) and IED(sp) density in the MD (left panel, positive correlation), and MD slow SP(pure) density (right panel, negative). Bottom panel shows the positive correlation between MD fast SP(pure) and seizure prevalence.*

*Supplementary Table 2. Overall absolute number, density, and mean duration of sleep spindles, separately for the ANT and MD averaged across each derivation within the same nucleus, separately for the spindle type (slow/fast).*

| | | **Absolute numbers** | | | | **Density** | | | | **Duration** | | | |
|---|---|---|---|---|---|---|---|---|---|---|---|---|---|
| | | ANT | | MDT | | ANT | | MDT | | ANT | | MDT | |
| | | Slow | Fast | Slow | Fast | Slow | Fast | Slow | Fast | Slow | Fast | Slow | Fast |
| Patients | #1 | 1594 | 1697 | 1748 | 1610 | 6.402 | 6.815 | 7.020 | 6.466 | 0.921 | 1.042 | 1.056 | 1.028 |
| | #2 | 2395 | 2540.5 | 2474 | 2431 | 7.589 | 8.050 | 7.839 | 7.703 | 1.187 | 1.058 | 1.186 | 0.961 |
| | #3 | 2035.5 | 1635 | 1921 | 1478 | 5.544 | 4.453 | 5.232 | 4.026 | 0.917 | 0.846 | 0.872 | 0.837 |
| | #4 | 2323 | 1139.67 | | | 8.623 | 4.230 | | | 1.277 | 0.867 | | |
| | #5 | 744 | 1257 | 745 | 1253 | 3.621 | 6.118 | 3.626 | 6.098 | 0.808 | 1.128 | 0.804 | 1.133 |
| | #6 | 1322.25 | 1464 | | | 6.329 | 7.007 | | | 0.972 | 1.047 | | |
| | #7 | 2448 | 2284 | 2532.5 | 2143 | 6.957 | 6.491 | 7.197 | 6.090 | 1.696 | 0.989 | 1.549 | 0.958 |
| | #8 | 1714.83 | 1172.67 | | | 7.268 | 4.970 | | | 1.055 | 0.849 | | |
| | #9 | 3102.33 | 1757 | 2909 | 1611 | 8.542 | 4.838 | 8.009 | 4.436 | 1.261 | 0.835 | 1.260 | 0.843 |
| | #10 | 2416 | 2352.67 | 2895.5 | 2348.5 | 7.385 | 7.192 | 8.851 | 7.179 | 1.025 | 1.176 | 1.079 | 1.168 |
| | #11 | 497.667 | 254.667 | | | 3.692 | 1.889 | | | 0.792 | 0.731 | | |
| | #12 | 1920 | 1429 | 1893 | 864.5 | 7.860 | 5.850 | 7.750 | 3.539 | 1.560 | 0.888 | 1.650 | 0.813 |
| | #13 | 1770.5 | 1052 | 1791 | 803 | 7.502 | 4.458 | 7.589 | 3.403 | 1.689 | 0.868 | 1.665 | 0.838 |
| | #14 | 2164 | 2695.75 | | | 5.659 | 7.050 | | | 0.914 | 1.036 | | |
| | #15 | 1004 | 849 | 986 | 907 | 7.545 | 6.380 | 7.410 | 6.816 | 1.549 | 0.962 | 1.611 | 1.028 |

*Supplementary Table 3. Absolute number and density of thalamic sleep spindles, averaged across each derivation within the same nucleus, separately for the spindle type (slow/fast) and association with IEDs and ripples (SP(pure/IED/ripple)).*

| | | **ANT** | | | | | | **MDT** | | | | | |
|---|---|---|---|---|---|---|---|---|---|---|---|---|---|
| | | Slow | | | Fast | | | Slow | | | Fast | | |
| | | pure | IED | ripple | pure | IED | ripple | pure | IED | ripple | pure | IED | ripple |
| Absolute numbers | #1 | 1086 | 362 | 146 | 1182 | 366 | 149 | 1349 | 287 | 112 | 1247 | 259 | 104 |
| | #2 | 1968 | 275.5 | 151.5 | 2214 | 198.5 | 128 | 2174 | 159 | 141 | 2194 | 136 | 101 |
| | #3 | 1475 | 316 | 244.5 | 1113 | 299 | 223 | 1496 | 292 | 133 | 1076 | 295 | 107 |
| | #4 | 1871.667 | 207.333 | 244 | 886.333 | 142.333 | 111 | | | | | | |
| | #5 | 471 | 234 | 39 | 906 | 263 | 88 | 467 | 234 | 44 | 869 | 271 | 113 |
| | #6 | 936.25 | 227 | 159 | 1007 | 268.75 | 188.25 | | | | | | |
| | #7 | 1659 | 526 | 263 | 1672 | 432 | 180 | 1920.5 | 311 | 301 | 1679.5 | 265 | 198.5 |
| | #8 | 1237 | 206.833 | 271 | 849.833 | 151.5 | 171.333 | | | | | | |
| | #9 | 1832 | 553 | 717.333 | 979.667 | 310.333 | 467 | 1684 | 334 | 891 | 893 | 222 | 496 |
| | #10 | 1404.333 | 859.667 | 152 | 1118.667 | 1077.667 | 156.333 | 1668.5 | 1052 | 175 | 921 | 1298.5 | 129 |
| | #11 | 300.667 | 134.333 | 62.667 | 142.333 | 71.667 | 40.667 | | | | | | |
| | #12 | 1230 | 468 | 222 | 910 | 384.5 | 134.5 | 1302.5 | 404 | 186.5 | 569 | 250 | 45.5 |
| | #13 | 1266.5 | 138.5 | 365.5 | 707 | 95.5 | 249.5 | 853 | 728 | 210 | 420 | 272 | 111 |
| | #14 | 1589 | 384 | 191 | 1978.5 | 465.25 | 252 | | | | | | |
| | #15 | 839 | 104 | 61 | 731 | 87 | 31 | 766 | 135 | 85 | 742 | 91 | 74 |
| Density | #1 | 4.361 | 1.454 | 0.586 | 4.747 | 1.470 | 0.598 | 5.418 | 1.153 | 0.450 | 5.008 | 1.040 | 0.418 |
| | #2 | 6.236 | 0.873 | 0.480 | 7.015 | 0.629 | 0.406 | 6.888 | 0.504 | 0.447 | 6.952 | 0.431 | 0.320 |
| | #3 | 4.018 | 0.861 | 0.666 | 3.032 | 0.814 | 0.607 | 4.075 | 0.795 | 0.362 | 2.931 | 0.804 | 0.291 |
| | #4 | 6.948 | 0.770 | 0.906 | 3.290 | 0.528 | 0.412 | | | | | | |
| | #5 | 2.292 | 1.139 | 0.190 | 4.409 | 1.280 | 0.428 | 2.273 | 1.139 | 0.214 | 4.229 | 1.319 | 0.550 |
| | #6 | 4.481 | 1.086 | 0.761 | 4.820 | 1.286 | 0.901 | | | | | | |
| | #7 | 4.715 | 1.495 | 0.747 | 4.752 | 1.228 | 0.512 | 5.458 | 0.884 | 0.855 | 4.773 | 0.753 | 0.564 |
| | #8 | 5.243 | 0.877 | 1.149 | 3.602 | 0.642 | 0.726 | | | | | | |
| | #9 | 5.044 | 1.523 | 1.975 | 2.697 | 0.854 | 1.286 | 4.637 | 0.920 | 2.453 | 2.459 | 0.611 | 1.366 |
| | #10 | 4.293 | 2.628 | 0.465 | 3.420 | 3.294 | 0.478 | 5.100 | 3.216 | 0.535 | 2.815 | 3.969 | 0.394 |
| | #11 | 2.230 | 0.997 | 0.465 | 1.056 | 0.532 | 0.302 | | | | | | |
| | #12 | 5.035 | 1.916 | 0.909 | 3.725 | 1.574 | 0.551 | 5.332 | 1.654 | 0.764 | 2.329 | 1.023 | 0.186 |
| | #13 | 5.367 | 0.587 | 1.549 | 2.996 | 0.405 | 1.057 | 3.614 | 3.085 | 0.890 | 1.780 | 1.153 | 0.470 |
| | #14 | 4.155 | 1.004 | 0.499 | 5.174 | 1.217 | 0.659 | | | | | | |
| | #15 | 6.305 | 0.782 | 0.458 | 5.493 | 0.654 | 0.233 | 5.757 | 1.015 | 0.639 | 5.576 | 0.684 | 0.556 |
